# Gradients of function between sensory drive and working memory in human frontal cortex

**DOI:** 10.64898/2026.08.25.747005

**Authors:** Thomas Possidente, Vaibhav Tripathi, Sangil Lee, David C. Somers

## Abstract

The coordination of sensory processing and working memory (WM) is fundamental to cognition. Spatial organization of sensory processing and WM is known to be broadly distributed across the cortex, but finer-scale organization at the interfaces between these functions remains understudied. Although the notion of sharp parcellations of cortex into distinct functional modules dominates the field, a growing body of works support graded changes in function and anatomy in some cortical zones. Based on this and potential advantages of gradient organizational structure in frontal cortex, we hypothesized that sensory-WM interfaces in the frontal cortex are gradient-like, not boundary-like. We examined twenty bilateral cortical regions that participate in visual/auditory WM tasks. In five frontal cortical regions, group-level WM activation overlapped with sensory drive, but was spatially shifted. We compared subject-level (N=20) boundary and gradient models of change in function. Strong individual-level evidence for sensory-WM gradients was observed in pre-supplementary motor area, ventral premotor cortex, and anterior insula in both modalities and in dorsal premotor cortex for visual WM. Conversely, dorsolateral pre-frontal cortex yielded mixed results, favored distinct WM and sensory regions in the left hemisphere, and gave some evidence for gradients in the right hemisphere. These results provide evidence that sensory and WM regions in frontal cortex are largely not distinct with sharp boundaries at their interfaces but instead bleed into each other to form local rostral-caudal sensory-WM gradients. We speculate these gradients may allow efficient interfacing between sensory and WM representations, and/or fine-grained, task-dependent shifting between bottom-up sensory and top-down influences.

**Significance Statement:** Cognitive neuroscience strongly adheres to the notion that the cerebral cortex is functionally parcellated into distinct regions with sharp anatomical boundaries. In contrast, modern neuroanatomical analysis indicates that, across cortex, boundaries vary from abrupt to gradual. Here, we examine the interface between sensory drive and working memory representations and observe meso-scale gradients (∼ 1 cm) in multiple regions of human frontal cortex. The observed functional gradients span regions previously defined as having sharp boundaries. Gradients offer potential benefits for working memory to balance faithful encoding of stimulus information with cognitive priorities necessary to support behavioral goals. The findings have important implications for understanding the neural architecture of working memory and more broadly for the functional parcellation of the cerebral cortex.

## Introduction

The coordination of sensory processing and working memory (WM) is fundamental to cognition. Sensory WM holds representations of remembered stimuli but also must distinguish these from incoming perceptual representations. The classic solution to this dual-representation problem is to hold mnemonic representations in frontal cortex and perceptual representations in posterior cortices (Constantinidis et al., 2001; Curtis & D’Esposito, 2003; Y. Xu, 2017). Although this is a useful general heuristic, it is complicated by findings showing that the contents of sensory WM are encoded in a distributed network spanning much of the brain (Christophel et al., 2017; Serences, 2016) and sensory stimuli drive activity exists within frontal cortex, even in the absence of a cognitive task in humans (Benson et al., 2018; Noyce et al., 2022; Saron et al., 2001) and non-human primates (Boussaoud & Wise, 1993; Funahashi et al., 1990; Romanski & Goldman-Rakic, 2002). Additionally, these two functions are often localized to similar cortical regions. These findings highlight the need for a finer scale examination of sensory drive and sensory WM-activated cortical regions. Here, we perform fMRI analyses (N=20) of visual and auditory sensory drive and WM, focusing on meso-scale functional organization in frontal cortices.

We consider three types of organizational structure that might occur in a cortical zone that exhibits activation for both sensory drive and sensory WM: 1) stimulus drive and WM activation patterns may fully overlap; 2) distinct sensory and WM representational zones may exist separately at a fine scale; 3) stimulus drive and WM representations may constitute a meso-scale functional gradient. The full overlap hypothesis appears highly plausible in early sensory cortices, given evidence in support of the sensory-recruitment hypothesis (Albers et al., 2013; Christophel et al., 2017; Pasternak & Greenlee, 2005).

The plausibility of the distinct subzones hypothesis is supported by fMRI studies employing spatially precise within-subject methods to reveal finer scale functional organization than previously assumed based on group-averaging methods (Braga & Buckner, 2017; Gratton et al., 2020; Michalka et al., 2015; Noyce et al., 2017). A leading cortical parcellation identifies over 50 distinct frontal regions (Glasser et al., 2016), and recent studies suggest even finer functional subdivisions (Assem et al., 2025; DiNicola & Buckner, 2026; Noyce et al., 2022). However, the vast complexity of cognitive functions embodied in frontal cortex may well exceed the plausible number of anatomically distinct domains, and there are plausible alternative accounts that deserve careful examination.

The meso-scale functional gradients hypothesis questions the assumption of hard boundaries between functional areas in the cortex that has dominated since the work of Brodmann (Brodmann, 1909) and continues today with modern functional brain parcellations (Buckner et al., 2011; Glasser et al., 2016; Schaefer et al., 2018). However, there is a long-held minority view that sharp cytoarchitectonic boundaries are the exception rather than the rule (Bailey & Von Bonin, 1951). Modern analysis indicates that many cortical architectonic boundaries are more gradual than abrupt (Amunts & Zilles, 2015). These ideas have only recently begun to be examined in functional neuroimaging, with a few fMRI studies demonstrating meso-scale functional connectivity gradients in cortex (Farrugia et al., 2024; Lefco et al., 2020; Shen et al., 2023; Tian & Zalesky, 2018). Here, we examine whether sensory-WM interfaces in frontal cortex may also be gradient-like, not boundary-like.

Our analysis begins by identifying cortical zones strongly driven by both sensory WM and by stimulus drive in the absence of a WM task. We then examine each of these cortical zones for evidence of spatial offsets in stimulus drive and WM activation at a group level and perform individual subject analyses directly comparing the subzone (boundary) and gradient hypotheses. The findings reveal positive evidence for the presence of meso-scale functional sensory-WM gradients for both visual and auditory activation in several frontal cortical areas.

## Methods

A separate analysis of the task fMRI data used here was previously published in Possidente, Tripathi et al. (2026). Duplicated text reporting on participants, task paradigm, and preprocessing is indicated in *italics*.

### Participants

*Twenty-four participants (aged 18 to 43; 11 men and 13 women [self-identified]) were recruited from the Boston University community. All procedures in this study were approved by the Institutional Review Board of Boston University (4928E). All ethical regulations relevant to human research participants were followed. Each participant gave written informed consent to participate in the study and received monetary compensation. All participants indicated that they had normal or corrected to normal vision and normal hearing. Two authors participated in the study (T. P., V. T.).* All subjects performed visual and auditory working memory and sensorimotor control tasks during fMRI for this study. Four participants were rejected from analysis - one for ROI identification problems, one for head movement, and two for lack of fixation condition – leaving twenty participants. Rejection criteria details are given below.

### Working Memory Paradigm

*All participants performed 4 tasks in the scanner: 2-back visual working memory, 2-back auditory working memory, visual sensorimotor control (SMC), and auditory sensorimotor control* (Figure 1)*. Stimuli display and task timing control for the 2-back WM tasks were performed using the Psychopy Toolbox (v2021.1.4;* www.psychopy.org (Peirce et al., 2019)*) in Python (v. 3.10* www.python.org*) running on a laptop running Windows 11 and displayed using a liquid crystal display projector that back-projected onto a screen placed within the scanner bore. Auditory stimuli were presented through MRI-safe in-ear earphones (Sensimetrics Model S14; Sensimetrics;* www.sens.com*). Stimuli were presented for 1 second each, with a 1 second gap between stimuli; only one modality was presented at a time. The visual stimuli were Gabor patches with spatial frequencies in the range of 1 to 6 cycles per degree. In the visual 2-back task, subjects judged whether the spatial frequency of the stimulus matched that presented two stimuli previously. After the presentation of the first two stimuli in a block, matches occurred on 50% of the trials. Gabor orientation varied randomly and was irrelevant to the match judgement. In the auditory 2-back task, the stimuli were auditory ‘warbles’ with either a 500 or 1000 Hz base pitch modulated by an oscillatory amplitude envelope with frequencies ranging from 3 Hz to 20 Hz. Subjects judged whether the envelope frequency matched that from two stimuli earlier. Pitch was varied randomly and was irrelevant to the match judgement. For both auditory and visual stimuli, the trials consisted of high frequency stimuli (i.e., odd numbered trials) alternated with low frequency trials (i.e., even numbered trials); this afforded the opportunity to make the matching task difficult while also presenting a broader range of stimuli. The percentage change in the modulatory frequency for non-match trials was adjusted on a trial-by-trial basis using independent staircasing procedures for visual and auditory stimuli (stepsize of 1%). Individual thresholds were identified for each subject in a behavioral session prior to scanning and were updated during the scans. The behavioral training session included presentation of recorded acoustic noise of the scanner to simulate the auditory distractions of the scanner*.

**Figure 1.**
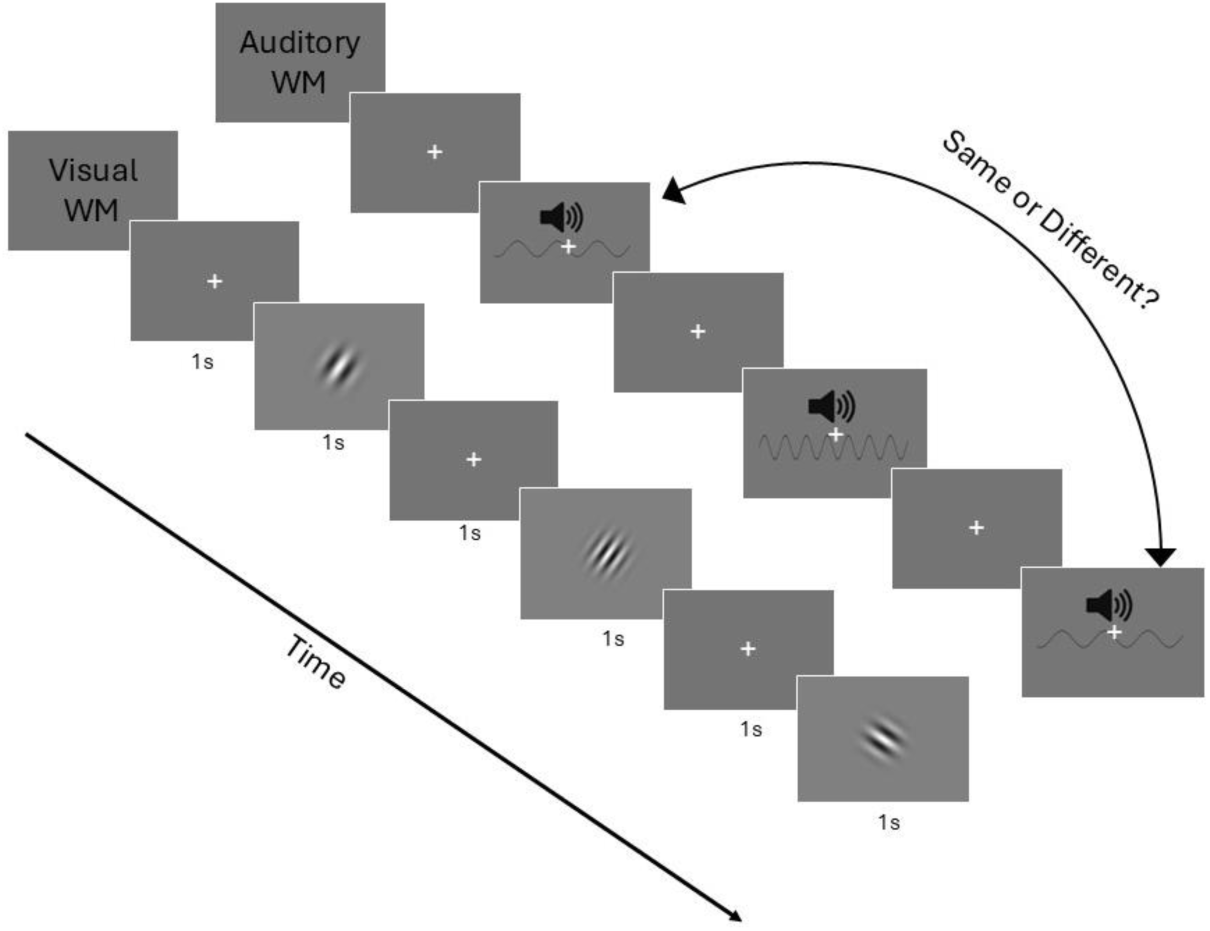
Visual and auditory 2-back WM task summaries. For the visual WM task, participants were asked whether the current Gabor filter spatial frequency matched the one presented two stimuli previously. For the auditory WM task, participants were asked whether the warble envelope volume modulation frequency matched the one presented two stimuli previously. Task performance required participants to maintain a high-precision representation of both the most recent even and odd-numbered trials. Figure reproduced with permission from Possidente, Tripathi et al. (2026).

*During the sensorimotor control tasks, auditory or visual stimuli were presented without 2-back repeats, and participants were instructed not to do the task but instead to arbitrarily press either of the two response buttons at the end of each trial. The relevant blocks in a single run consisted of 2 blocks of visual WM, 2 blocks of auditory WM, 1 block of visual sensorimotor control, 1 block of auditory sensorimotor control, and 1 fixation condition block without stimuli or responses. Block order was randomized from run to run for all participants. Each block lasted 32s and contained 16 trials. Participants were given feedback indicating correct and incorrect trials. During the sensorimotor control conditions, feedback was given if they had pressed the key or not.* All participants had 4 runs.

### MR Imaging

*All scans were performed at Boston University’s Cognitive Neuroimaging Center on a Siemens 3-Tesla Prisma scanner with a 64-channel head coil. Structural images were collected for each participant with T1-weighted high-resolution magnetization-prepared rapid gradient multi-echo sequence (MP-RAGE,* (van der Kouwe et al., 2008)*) (1mm isotropic voxels, 176 slices, repetition time [TR] = 2530ms, echo times [TE] = 1.69, 3.55, 5.41 and 7.27ms, flip angle [FA] = 7°, field of view [FOV] = 256mm, GRAPPA [iPAT* (Griswold et al., 2002)*] acceleration=3, TA=4min 23s). All blood-oxygen-level-dependent (BOLD) data were collected via T2*-weighted echo-planar imaging (EPI) pulse sequence that employed multiband RF pulses and simultaneous multi-slice (SMS) acquisition* (Cauley et al., 2014; Feinberg et al., 2010; Moeller et al., 2010; Setsompop et al., 2012; J. Xu et al., 2013)*. Task fMRI (69 slices, TR=2000ms, TE=35ms, FA=80°, 2.2mm isotropic voxels, FOV=207mm, SMS=3, TA=7min 46s) was also acquired. Pairs of reversed phase-encoded (anterior-posterior and posterior-anterior) spin echo field maps were also acquired for subsequent EPI de-warping*.

### MRI Preprocessing

*Structural and functional data preprocessing and general linear modeling were performed using Freesurfer (version 7.4.1, available at* http://surfer.nmr.mgh.harvard.edu/) (Fischl, 2012)*. Anatomical reconstruction (recon-all) and registration (register-sess) were performed* (Dale et al., 1999; Fischl et al., 2002) *with human quality control inspection. Preprocessing included removal of non-brain tissue using a hybrid watershed/surface deformation procedure* (Ségonne et al., 2004)*, automated Talairach transformation, intensity normalization* (Sled et al., 1998)*, tessellation of the gray matter white matter boundary, automated topology correction* (Fischl et al., 2001; Segonne et al., 2007)*, and surface deformation following intensity gradients to optimally place the gray/white and gray/cerebrospinal fluid borders at the location where the greatest shift in intensity defined the transition to other tissue* (Dale et al., 1999) *(mkbrainmask-sess). Functional data preprocessing included motion correction (mc-sess), FSL’s “topup” and “applytopup” functions* (Jenkinson et al., 2012; Smith et al., 2004) *for field map distortion correction, slice-time correction (stc-sess), and individual volume to fsaverage surface transformation (163,842 vertices, rawfunc2surf-sess).* For probabilistic group level analysis and individual subject ROI identification, 5mm full width at half max (FWHM) Gaussian kernels were used to smooth data. All individual subject-based analyses (contrast gradient visualization and boundary vs. gradient modeling) used unsmoothed data. *General linear modeling included one regressor per stimulus condition convolved with the SPM canonical hemodynamic response function, as well as a nuisance regressor for linear drift removal, and 12 motion correction regressors per run (3 translation, 3 rotation, and their derivatives)*.

*Runs were rejected if there was more than 1mm average head displacement from midpoint position during the run (calculated using Freesurfer’s “mc-sess”). One full subject was rejected as a result.* One additional participant was rejected from analysis due to overall low activation at a liberal p<.05 uncorrected threshold (ex. no significant activity in the auditory cortex). Lastly, two participants were rejected because they did not complete the fixation condition of the task, a condition necessary for computing sensory drive contrasts. This left 20 participants for analysis.

### Overview of Gradient Analysis Workflow

To examine whether meso-scale functional gradients exist between sensory drive and WM representations in cortex, we took a multistep approach that began with coarse group-level analyses across the cortex and ended with fine within-subject analyses in candidate ROIs that passed several criteria. This workflow is summarized in Figure 2. Briefly, we first identified an initial set of 20 large, bilateral ROIs for vision and/or audition which exhibited strong responses to sensory drive and/or WM. We narrowed this set to those ROIs that exhibited strong responses to both sensory drive and WM, and further narrowed this to a final set of candidate ROIs that exhibited a shift in the center of mass of the sensory drive and WM activation patterns within the ROI at the group level. The particular thresholds for these ROI rejection criteria were determined prior to analysis through an analysis plan document for a related project (https://doi.org/10.17605/OSF.IO/3TBXC). We then examined individual subject patterns within these ROIs testing them against gradient and sharp boundary models. These steps are detailed below.

**Figure 2.**
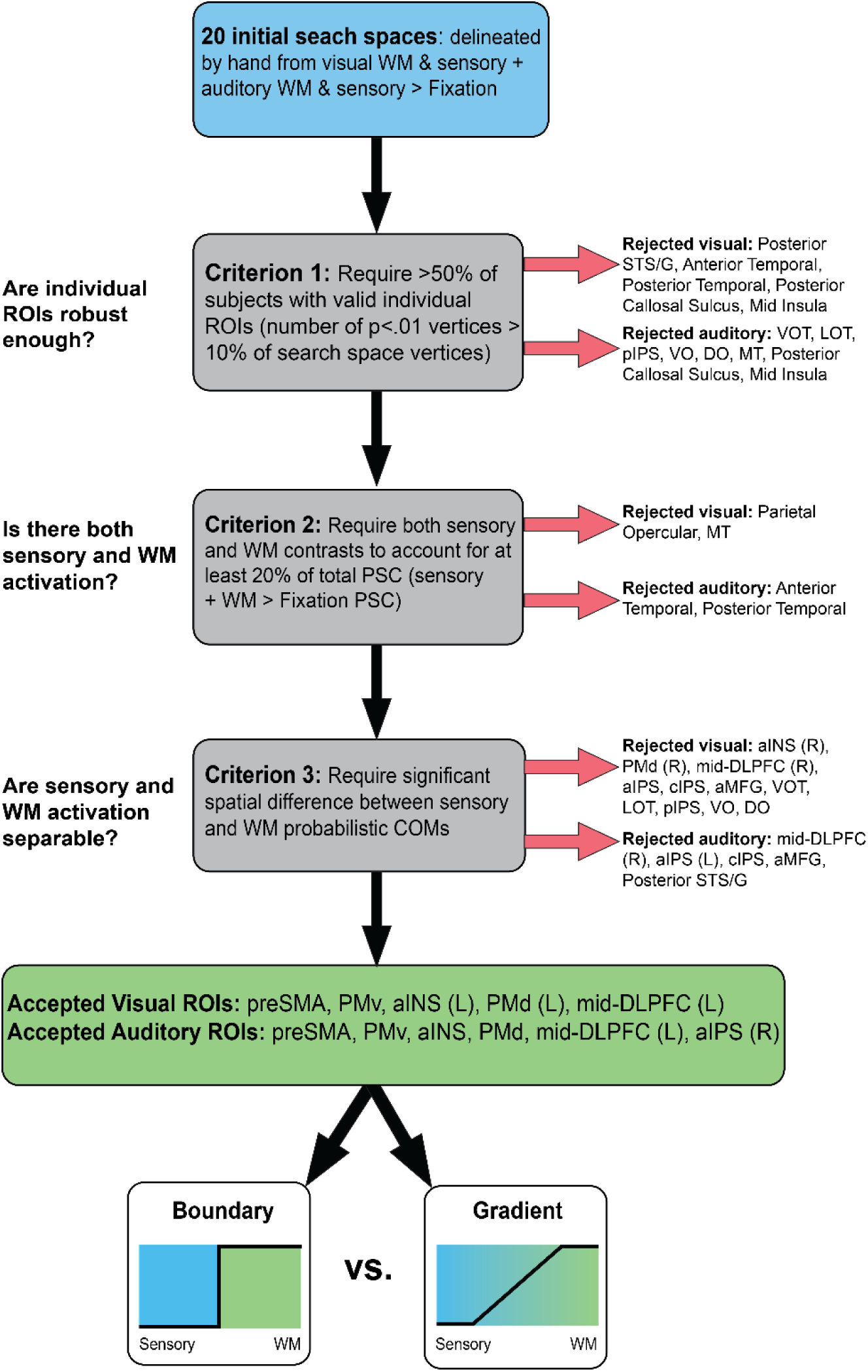
ROI rejection criteria flowchart.

### ROI Identification

Individual ROIs were created using a group-constrained subject-specific approach. To create the probabilistic search spaces used to constrain individual subject ROI identification, we first computed p<.05 uncorrected whole-cortex masks based on the contrast of all visual conditions (WM and sensorimotor control [SMC]) against fixation and all auditory conditions against fixation. We then added the subject-level masks together and thresholded the resulting map at 5 subjects (25%). We then took the union of the resulting auditory and visual probabilistic maps. The resulting union map represents areas of cortex that are likely to respond to visual or auditory sensory drive or WM. Two authors (T.P. and D.S.) delineated probabilistic ROI search spaces by hand on the union map using hotspots and restriction points in the maps, functional areas from literature, and anatomical landmarks (Figure 3).

**Figure 3.**
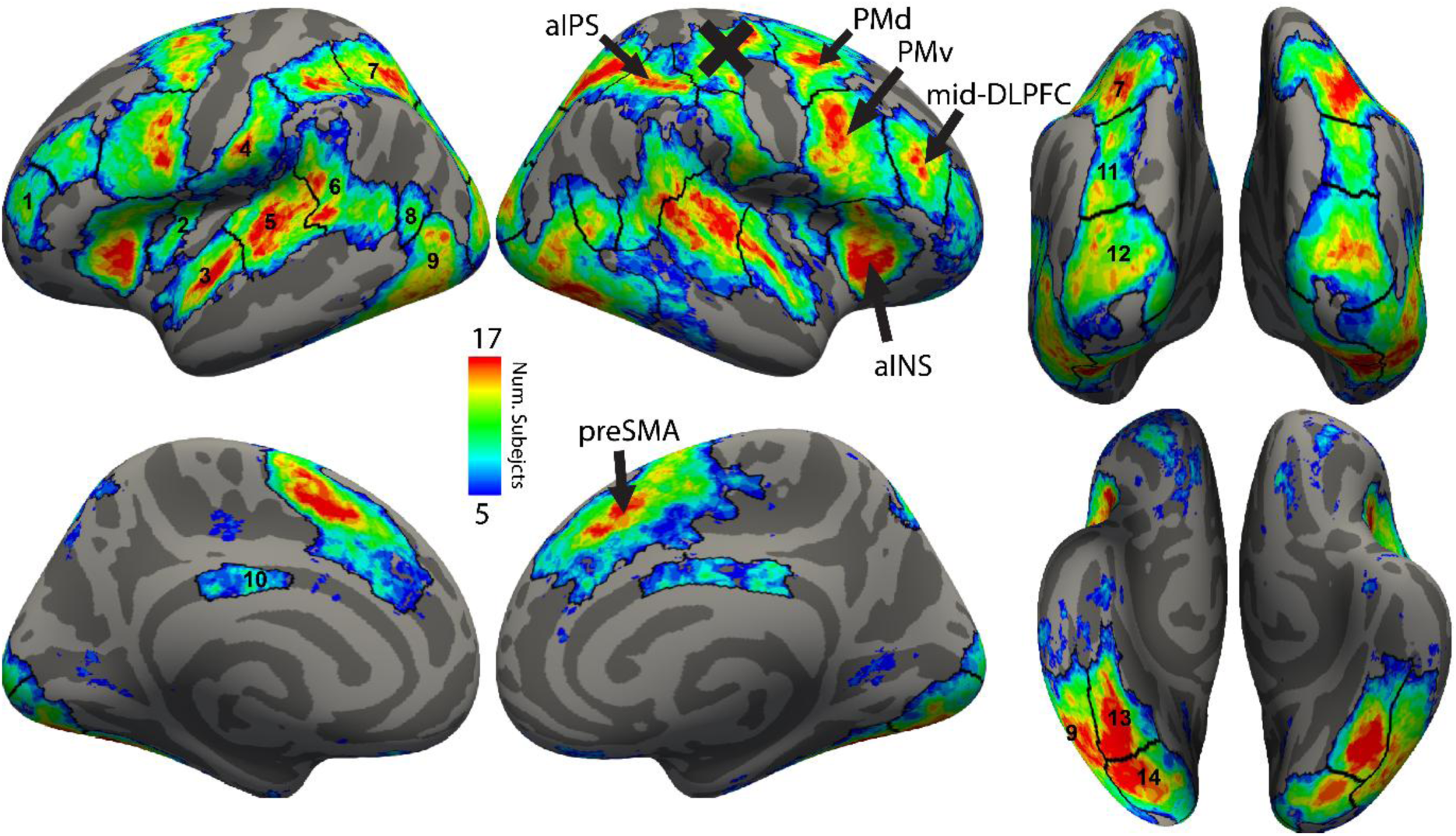
Lateral, medial, posterior, and ventral views of probabilistic activation maps for the union of all visual conditions against fixation and all auditory conditions against fixation for both hemispheres. ROI delineations are shown as black lines. Sensory-WM interface candidate ROIs are labeled with arrows. A motor cortex ROI on the right hemisphere is X’d out to indicate that it is a motor confound (left hand button press) and does not indicate sensory or WM activation. Numbered ROIs were rejected from gradient analysis and are as follows: 1. Anterior middle frontal gyrus (aMFG), 2. Middle insula, 3. Anterior temporal, 4. Parietal opercular, 5. Posterior temporal, 6. Posterior superior temporal sulcus/gyrus (pSTS/G), 7. Central intraparietal sulcus (cIPS), 8. Middle temporal visual area (MT), 9. Lateral occipital temporal (LOT), 10. Posterior callosal sulcus, 11. Posterior intraparietal sulcus (pIPS), 12. Dorsal occipital (DO), 13. Ventral occipital temporal (VOT), 14. Ventral occipital (VO).

Individual subject visual and auditory ROIs were then automatically identified using the following contrasts respectively: visual WM & visual SMC against fixation, and auditory WM & auditory SMC against fixation. For each contrast separately, individual subject ROIs were created from p<.01 uncorrected vertices within each probabilistic ROI search space. Individual ROIs with fewer than 10% of the number of vertices in the corresponding probabilistic search space were rejected. ROIs that were present in fewer than 50% of subjects were fully rejected from the analysis (Criterion 1 in Figure 2).

### Group-level Analysis of Probabilistic ROI Shifts

To identify candidate ROIs likely to contain interfaces between sensory drive and WM, we sought to reject ROIs that did not contain both sensory drive and WM activation. Thus, we required that the mean percent signal change (PSC) within each ROI across subjects for the WM contrast (WM against SMC) and sensory drive contrast (SMC against fixation), both be at least 20% of the full WM & SMC against fixation contrast. We did this for the visual and auditory contrasts separately (Criterion 2 in Figure 2).

We further sought to limit candidate ROIs to those with some spatial separation between sensory drive and WM activation (Criterion 3 in Figure 2). To determine significant spatial separation, we first calculated the probabilistic versions of the sensory drive and WM contrasts (vertices with p<.05, thresholded at 5 subjects). We then cut out and flattened each of the probabilistic ROI search spaces (using Freesurfer’s label2patch and mris_flatten) and calculated the center of mass (COM) on the cortex for each. COMs were calculated by weighing each vertex’s 2D coordinates by its probabilistic value (the number of subjects with significant activation in that vertex) and normalizing by the sum of all vertices’ probabilistic values. We then took the Euclidean distance between WM and sensory drive contrasts’ COMs as our statistic of interest. To test significant spatial differences between COMs, we repeated this process 10k times, permuting the individual subjects’ WM and sensory drive contrast labels randomly each time. We did this for visual and auditory probabilistic maps separately. The COM distances calculated from the permuted probabilistic maps in each ROI search space formed null distributions to compare against the original COM distances at p<.05 with Benjamini-Hochberg False Discovery Rate (BH-FDR) correction for all comparisons. After all rejection criteria, only the preSMA, PMv, aIns, PMd, aIPS, and mid-DLPFC remained for further subject-level analysis.

In order to estimate the group-level axis of greatest change from sensory to WM function in each ROI, we first took the group level sensory and WM contrast T-statistics in each flattened probabilistic ROI and subtracted them from each other at each vertex. We then took the weighted average of the 2D coordinates with positive T-statistics (indicating greater WM activation) and negative T-statistics (indicating greater sensory drive activation) and connected these points to form a group-level axis of greatest change for each ROI. These axes were used as starting estimates for individual subjects’ axes of greatest change during modeling.

### Individual Contrast Gradient Analysis

To gain a qualitative understanding of the WM and sensory drive spatial distributions at the individual subject-level, we plotted the difference between the WM and sensory contrasts unsmoothed T-statistics across the axis of greatest change for each ROI. To do this, we took each individual ROI and determined the axis across which there was the best division (boundary) between WM and sensory drive activity (see next section for details), then collapsed the difference T-statistic data from the ROI onto that axis. We then averaged T-statistics in 2.2mm bins and aligned all individual ROIs at their boundary point on the X-axis before plotting.

### Individual Gradient vs. Boundary Model Comparison

To model the structure of functional change across individuals’ ROIs, we fit step (boundary) and hinge (gradient) functions to individual T-statistic data on flattened cortical surface patches for each subject and ROI. To do this, we first capped the negative sensory and WM contrast T-statistics at zero then subtracted them from each other at each vertex. Capping the negatives at zero served to preserve the interpretability of the subtracted T-statistic maps and impacted only 11.95% of vertices on average for the candidate ROIs. Then, to reduce outliers we clipped T-statistics to 3 standard deviations from the median. We then rotated the individual ROI to align with the *group level axis* of greatest change. To more accurately fit each individual ROI, we rotated the ROI ±60° from the group-level axis in 1° increments, fitting the boundary model each time and taking the angle that resulted in the best r^2^ fit as the *individualized axis* of greatest change. For this process we collapsed the 2D difference T-statistic maps onto a single axis and fit the following nonlinear step function to the 1D data:

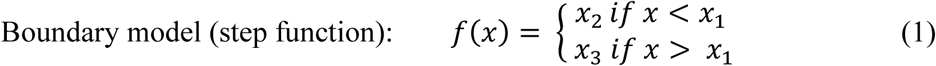

We used the boundary model for this process to steel-man the boundary hypothesis. Thus, we were then ready to compare the gradient and boundary model fits on the individual subject ROI along the axis that best separated the sensory and WM activation.

Next, to fit 1D gradient and boundary models to flattened 2D individual ROIs, we partitioned the ROIs into 2.2mm strips, parallel to the individualized axis of greatest change, and fit the boundary (equation 1) and gradient (equation 2) functions to each strip independently (Figure 4). Although more computationally intense, partitioning the ROI into thin strips mitigates the reduction in fit quality that would result from collapsing the full 2D data into 1D, where Y-axis variance could be expressed as noise around a potential step function. In contrast, during the individualized axis-finding process, we opted to collapse the full 2D data into 1D to avoid the greater computational burden of fitting each slice of each ROI for each subject at each potential angle (which would increase the number of models fitted by 120x). For each ROI, the mean r^2^, RMSE, and SSE across all strips were weighted by the number of vertices in each strip. RMSE and SSE were used to compute each model’s Bayesian information criterion (BIC) for model comparison. Strips with fewer than 20 data points were rejected.

**Figure 4.**
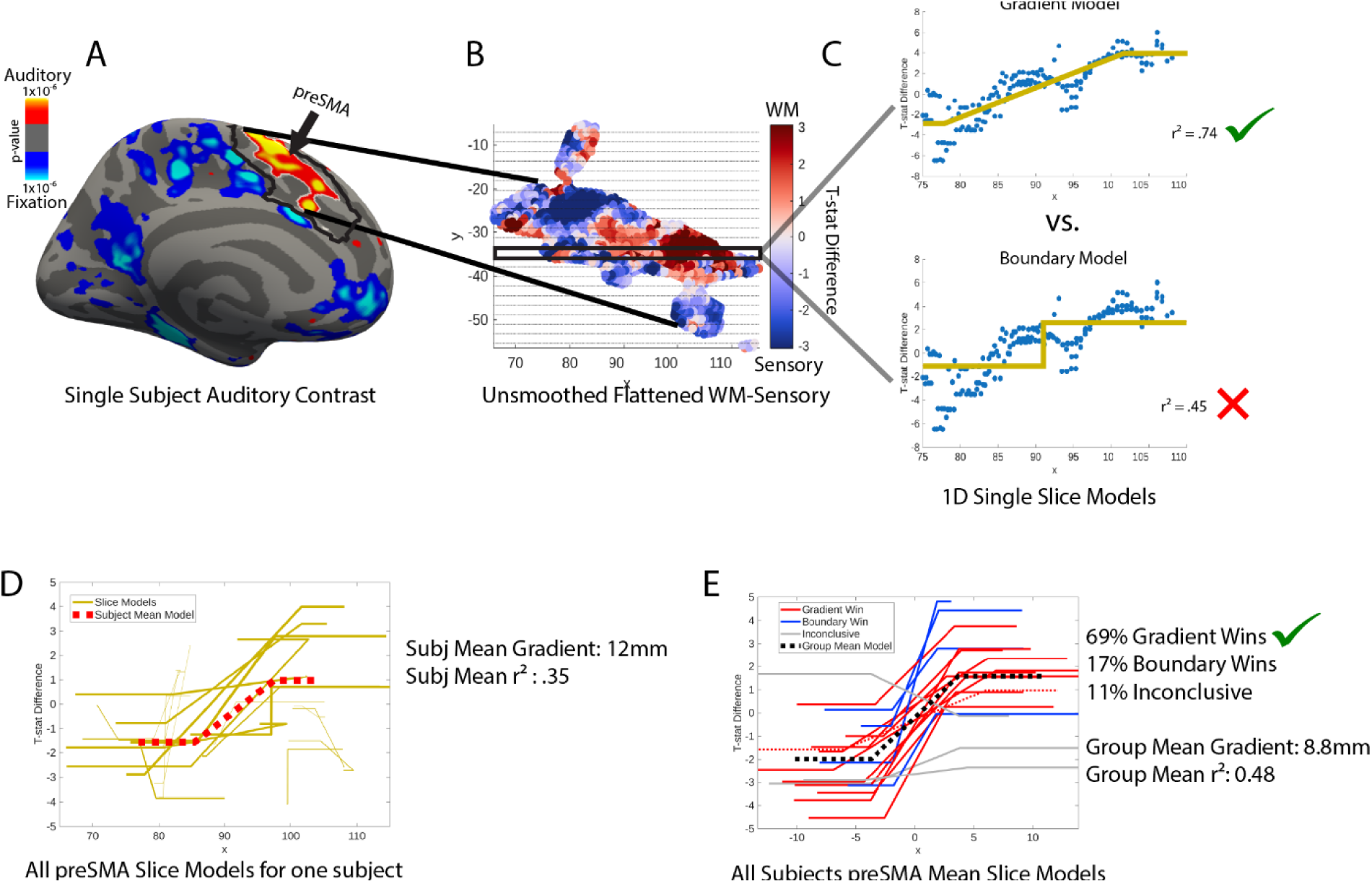
Process diagram for individual subject ROI gradient vs. boundary model comparison. A) Individual subject p-value map of all visual conditions (WM and SMC) against fixation. The black outline represents the probabilistic search space for the preSMA ROI while the white outline represents the p<.01 ROI inside the search space for this subject. B) The ROI from A is flattened and the difference of the unsmoothed T-statistics for the WM against SMC and SMC against fixation contrasts are projected. The X-axis is aligned with the axis of greatest change. Gray horizontal gridlines form slices that are 2.2mm in height. C) Data from a single slice is collapsed into 1D, then gradient and boundary models are fit to the data. This is repeated for each slice. D) Each slice’s hinge model is shown with line thickness corresponding to the number of data points in the slice. The red dotted line is the mean hinge model across all slices, weighted by the number of data points in each slice. This process is repeated for each subject. E) Each subject’s mean model is plotted, with gradient wins in red, boundary wins in blue, and inconclusive models in gray. All models’ x-values are aligned so that 0 corresponds to the best fit boundary point between sensory and WM activation for that subject/ROI. The dotted black line is the group mean model and the dotted red line is the subject mean model shown in panel D.

The following equation was used to fit the gradient model:

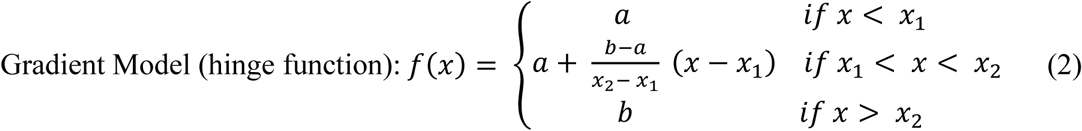

Because the boundary and gradient models had different numbers of parameters (3 and 4 respectively), we used BIC to adjudicate between models and required a BIC difference of 10, indicating strong evidence (Raftery, 1995), to rule in favor of one model over another. We also required an r^2^ > .1 to ensure the winning model fit the data to a reasonable level. Additionally, if the gradient model was preferred, but the gradient component of the fitted model was under 4.8mm, we considered it a win for the boundary model. This threshold was chosen by simulating randomly oriented functional boundary data in 2D, convolving with a 3.5mm FWHM Gaussian to account for the inherent point spread function (PSF) in fMRI, and down-sampling to 2.2mm voxels to simulate ground truth functional boundaries collected at fMRI resolution. Then, to mirror the modeling process used for the empirical data, we collapsed the simulated data into 1D perpendicular to the boundary and fit the hinge function to the 1D data. We repeated this process 1k times and took the 95^th^ percentile hinge model gradient length as our gradient length threshold. The 3.5mm Gaussian was chosen based on PSF estimation from Engel et al. (1997) for 1.5T, and is likely an overestimate for 3T. A summary of the requirements for boundary and gradient evidence is given in Table 1. This model comparison process was run for each ROI for visual and auditory contrast data separately.

**Table 1.** Shows the conditions under which the boundary or gradient model “wins”. All conditions not listed were considered inconclusive.

| BIC $\Delta > 10$<br>favoring step | BIC $\Delta > 10$<br>favoring hinge | $r^2 > .1$ | Gradient $> 4.8\text{mm}$ | Result |
| --- | --- | --- | --- | --- |
| ✓ | ✗ | ✓ | - | Boundary Evidence |
| ✗ | ✓ | ✓ | ✗ | Boundary Evidence |
| ✗ | ✓ | ✓ | ✓ | Gradient Evidence |

All models were fit using Matlab’s “fit” function with nonlinear least squares cost function using the trust region optimizer. For the gradient model, to ensure *x*_1_< *x*_2_, *x*_2_ was reparametrized to (*x*_1_ + *x*_2_) in equation 2 during model fitting. Parameters were initialized using the group-level T-statistic difference data (see Supplemental Table 1). Models in which the best fit was in the opposite direction expected by the group-level axis of greatest change were considered inconclusive (8.5% of models for preSMA, PMv, and aIns). All models converged successfully, with no optimization errors or warnings and parameters falling within reasonable ranges.

R-squared and gradient length for each ROI and modality were tested for significance using linear mixed effects models (LMMs) where hemisphere and subject were random effects and the model intercept term was used to test significance (no fixed effects). LMMs were chosen to account for repeated measures (left and right hemisphere from the same subject) and to easily deal with unequal missing data due to individual subject ROI rejections and whole hemisphere rejections for some ROIs. The model specifications in Wilkinson notation formatted for Matlab are as follows:

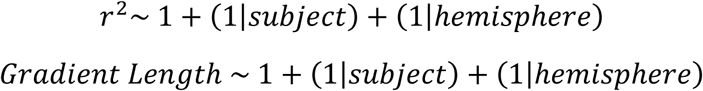

For ROIs where only one hemisphere was analyzed, the intercept was the only term in the model. R-squared was required to be significantly above .1 while gradient length was required to be significantly above 4.8mm. Similarly, 4.8 was subtracted from all gradient lengths prior to modeling. Each model’s intercept was tested at p<.05 correcting for 24 comparisons (6 ROIs, 2 modalities, 2 models each) using the BH-FDR procedure.

### Dice Coefficient Analysis

In order to determine how much the sensory and WM probabilistic maps overlapped with the core domain general parcels from the Human Connectome Project Multi-Modal Parcellation 1.0 (MMP), we performed Sørensen-Dice coefficient spatial overlap analysis (Dice, 1945; Sørensen, 1948) between the WM and sensory probabilistic maps for each modality and the core domain general MMP parcels in frontal cortex (8BM, 8C, IFJp, p9-46v, a9-46v, i6-8, AVI). This metric quantifies spatial similarity by taking 2 times the number of overlapped vertices and dividing by the sum of the vertices in each map:

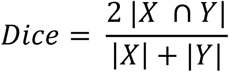

where |X| indicates number of elements in set X.

## Results

Subjects (N=20) performed 2-back visual working memory (WM) and 2-back auditory working memory tasks during fMRI (see Figure 1; Methods). Stimuli and task demands were chosen to require high fidelity WM of stimulus features (see Methods). Task accuracy did not differ significantly between visual and auditory WM tasks (visual WM=67.19% auditory WM=67.90%, t(19)=-.64, p=.53). All subjects also completed visual and auditory sensorimotor control (SMC) conditions where stimuli were presented and random button press responses were made on each trial, but no WM task was performed. WM activation was assessed through the contrast of WM and SMC conditions whereas sensory drive activation was assessed through the contrast of SMC and Fixation conditions.

### Group-level evidence for offset overlapping sensory and working memory frontal cortex regions

We implemented a multistep procedure to identify candidate gradient ROIs from group probabilistic activation maps and to test for gradients in individual subjects (see Figure 2; Methods). Briefly, we first constructed probabilistic cortical maps for the activation of sensory drive and/or WM for each modality and from this identified a candidate set of 20 ROIs per hemisphere (see Figure 3). We then excluded ROIs that failed to produce both robust sensory drive and robust WM activation. This process was carried out for visual and auditory probabilistic maps separately. We further limited our search for sensory-WM gradients to ROIs with significant shifts in the spatial pattern of the group probabilistic maps for sensory drive and WM. For each ROI, we tested whether the spatial center of mass (COM) of sensory drive activation (SMC vs. fixation) and WM activation (WM vs. SMC) were significantly spatially separated using a permutation test on probabilistic activation maps, with BH-FDR correction across ROIs and modality (visual, auditory). Only 6 ROIs (1 parietal, 5 frontal) showed significant spatial separation between sensory drive and WM activation at the group probabilistic level: pre-supplementary motor area (preSMA), dorsal premotor cortex (PMd), anterior insula (aIns), ventral premotor cortex (PMv), middle dorsolateral prefrontal cortex (mid-DLPFC), and anterior intraparietal sulcus (aIPS) (Table 2). These shifts between sensory drive and WM are illustrated in Figure 5 and Supplemental Figure 1. A rostral-to-caudal shift direction in the sensory drive to WM probabilistic maps was observed within several frontal regions, echoing the direction of the macroscale gradient reported in prior studies of cognitive control in the frontal lobe. However, mid-DLPFC shifted in the opposite direction.

**Figure 5.**
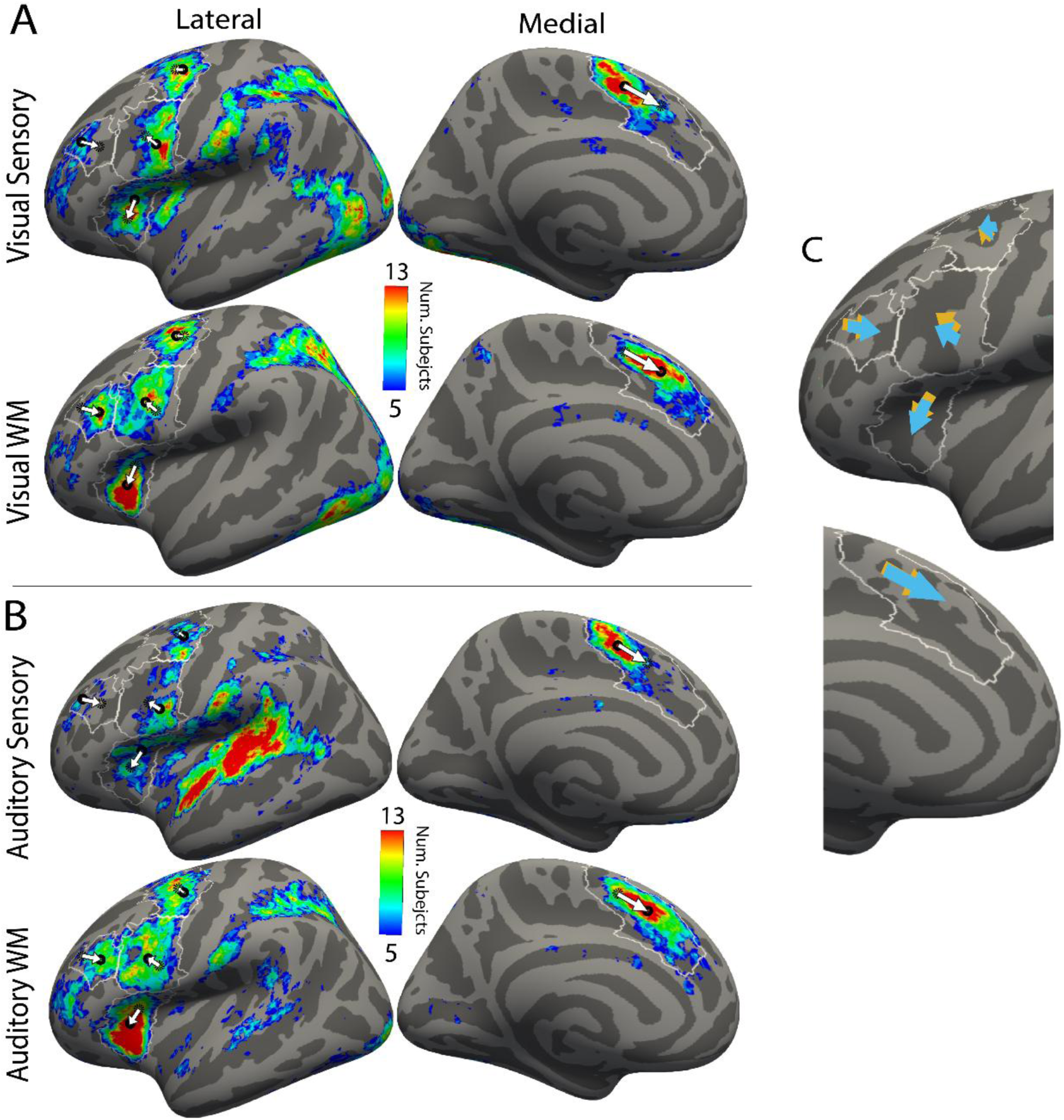
A) Probabilistic maps of visual sensory activation (top) and visual WM (bottom) with lateral (left) and medial (right) views. White arrows represent change of COM location from sensory to WM within each ROI. Black circles represent COMs for the map being displayed while black dotted circles represent COMs for the opposite gradient contrast. White borders show the probabilistic search spaces for each ROI. B) Same as A except using auditory sensory activation (top) and auditory WM (bottom). C) Visual (blue) and auditory (orange) sensory-WM gradient directions consolidated from A and B and enlarged. Right hemisphere results are shown in Supplemental Figure 1.

**Table 2.** Distance (mm) between sensory and WM probabilistic ROI centers of mass for each ROI. Green cells indicate significance under p<.05 BH FDR-corrected permutation testing. ROIs with no significant differences were excluded from the table.

| ROI | Hemisphere | Visual COM Distance (mm) | Auditory COM Distance (mm) |
| --- | --- | --- | --- |
| preSMA | L | **16.7** | **15.5** |
|  | R | **23.7** | **24.7** |
| PMv | L | **11.5** | **10.3** |
|  | R | **9.1** | **11.2** |
| aIns | L | **13.1** | **12.8** |
|  | R | 4.3 | **5.5** |
| PMd | L | **4.1** | **7.9** |
|  | R | 2.0 | **9.8** |
| mid-DLPFC | L | **11.8** | **13.5** |
|  | R | 4.7 | 3.3 |
| aIPS | L | 2.0 | 3.5 |
|  | R | 1.7 | **7.7** |

Thus far, we have demonstrated a set of ROIs that exhibit overlapping but shifted probabilistic maps for sensory drive and WM; however, this is insufficient to demonstrate a gradient structure. This sensory-WM interface structure suggests two potential explanations at the individual level: 1) separate homogeneous sensory drive and WM zones with individual heterogeneity in cortical boundaries resulting in overlap at the probabilistic level, or 2) a gradient between sensory drive and WM activation. To adjudicate between these possibilities, individual subject-level analysis was necessary.

### Individual subject-level ROI identification

Subject-level ROIs were identified using a group-constrained subject-specific procedure (see Methods) separately for visual and auditory contrasts. Only preSMA, PMv, aIns, PMd, aIPS, and mid-DLPFC ROIs were used for individual level analysis. Across modalities, hemispheres, and ROIs, 82.1% (279/340) of subject-level ROIs were successfully identified (see Supplemental Table 2 for detailed breakdown). These results are in line with previous individual-subject ROI identification rates from our lab (Possidente, Tripathi et al., 2026).

### Subject-level visualization reveals graded activation in sensory and working memory contrasts

These patterns of overlapped, shifted sensory drive and WM activations patterns were observed in individual subjects as well (see Figure 6, Supplemental Figure 2). To better understand the subject-level structure of change between WM and sensory drive across each ROI, we projected the WM-sensory T-statistic differences for individual subjects onto the axis of greatest contrast between WM and sensory drive activation, binned at 2.2mm (1 voxel length), and plotted the resulting spatial distributions for each subject aligned on the X-axis to the individualized best-fit boundary point between WM and sensory contrasts (see Methods, Figure 4). The resulting visualization of each contrast across each ROI (Figure 7 and Supplemental Figure 3) shows qualitative evidence for relatively gradual changes in WM vs. sensory drive across preSMA, PMv, and aIns in both modalities, with PMd showing noisier distributions and mid-DLPFC showing somewhat sharper changes in activation across a shorter span of cortex. Additionally, visualizing each contrast’s T-statistics separately (Supplemental Figures 4, 5) indicates that both contrasts are contributing to the overall trends in the difference T-statistics shown here (i.e. it is not the case that one contrast is constant while the other changes). Although there is heterogeneity in the individual distributions, preSMA, PMv, and aIns largely indicate that graded changes across the mean distribution are not caused by spatial variability in location of individual boundary-like step functions.

**Figure 6.**
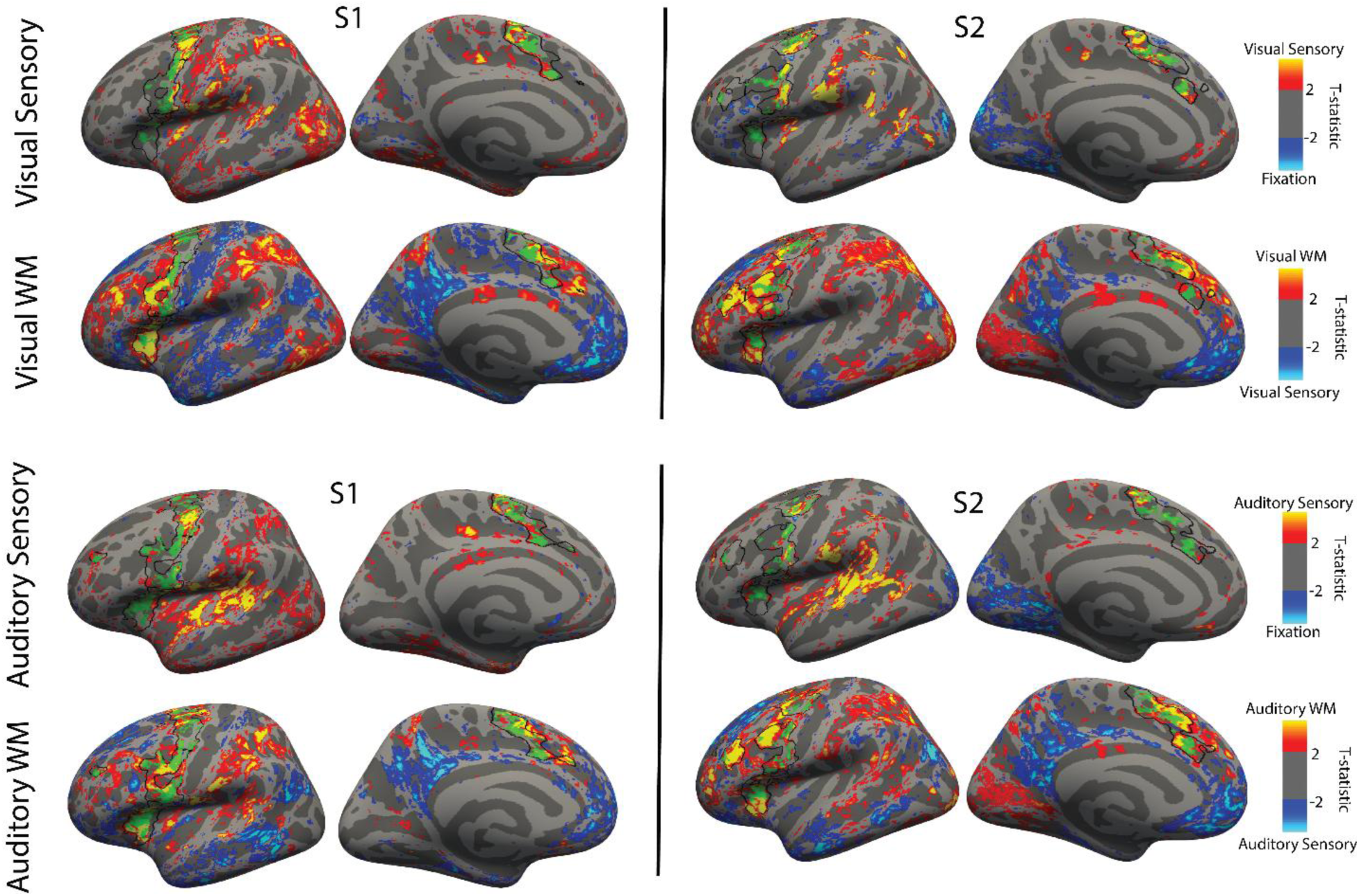
Arrangement is analogous to Figure 5A,B, but instead of probabilistic maps, two example subjects’ T-statistic maps are shown using a T>2 threshold. Black outlines show individual-specific ROIs (identified using WM and sensory vs fixation contrast for each modality separately). ROIs that did not pass the inclusion criteria are not outlined in black. Green patches represent areas of overlap between the WM and sensory T-statistic maps. The right hemisphere version of this figure is shown in Supplemental Figure 2.

**Figure 7.**
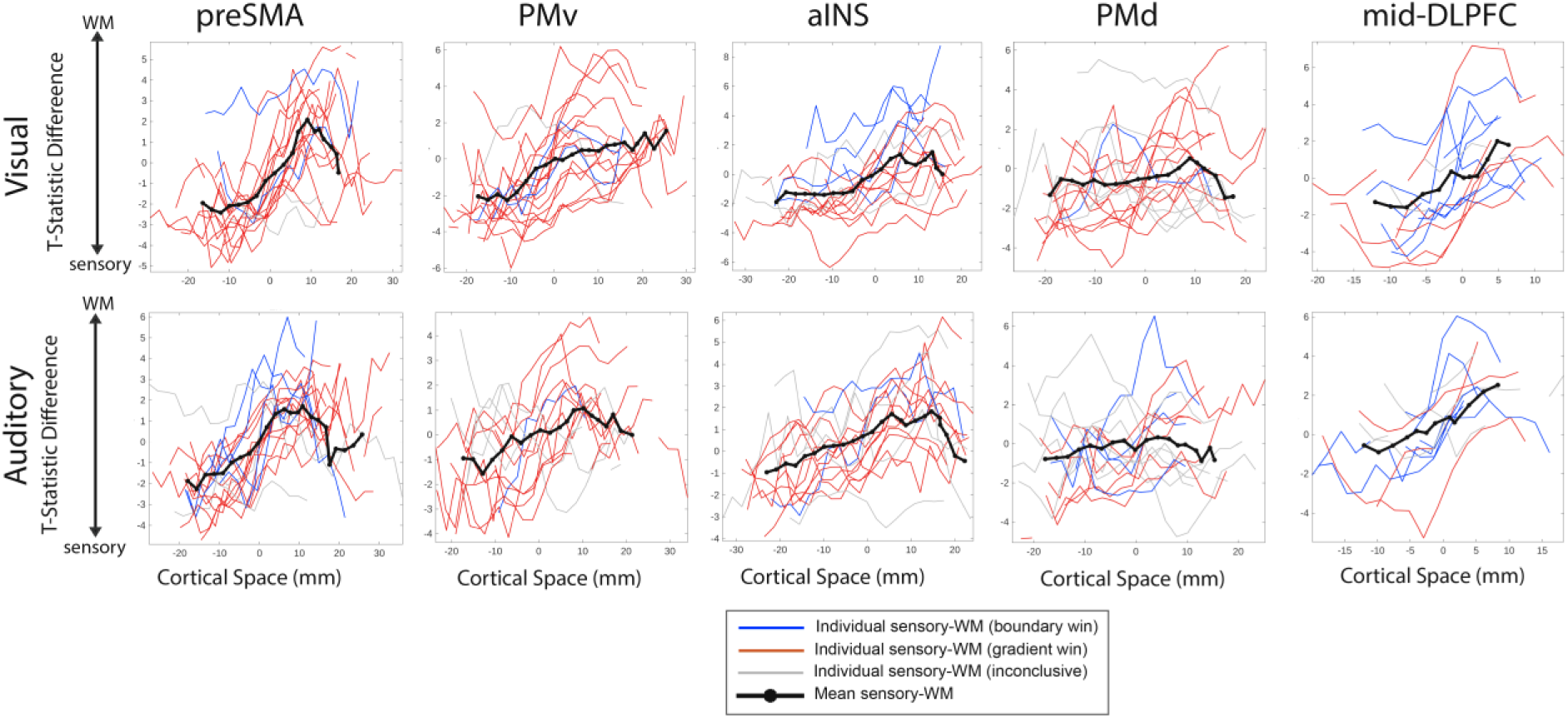
Binned T-statistics difference plotted across each ROI for each individual subject (visual and auditory). Thick black lines show the means of sensory/WM contrasts across subjects. Mean lines are only displayed if there are at least 5 subjects to average across. Each subject’s X-values are aligned so that 0 corresponds to the best fit boundary point between sensory and WM activation for that subject/ROI. Left hemisphere results are shown here (see Supplemental Figure 3 for right hemisphere).

Although this visualization provides some evidence in favor of the gradient hypothesis for preSMA, PMv, and aIns, it is limited by two interacting factors: 1) individual distributions are effectively smoothed due to spatial binning, which could obscure sharper changes in function, 2) gradient-like and boundary-like properties are not rigorously quantified at the individual subject level, leading to the possibility that some individual distributions are best described as boundary-like. Thus, we sought to address these limitations by explicitly quantitatively comparing boundary and gradient models in individual subjects.

### Quantifying individual subject-level evidence for sensory-working memory gradients

To directly compare boundary and gradient hypotheses for the interface regions identified at the group probabilistic level, we pitted step function (boundary) and hinge function (gradient) models of functional change against each other in subject-level ROIs. WM and sensory drive T-statistics were subtracted from each other at each vertex to form a flattened 2D surface of difference T-statistics. Rather than collapsing the full ROI patch of 2D data into 1D, models were fit independently to data in 2.2mm strips parallel to the axis of greatest change and fit statistics were aggregated across strips (Figure 4). Model selection used BIC with a difference threshold of 10 (strong evidence) and required an r-squared of more than .1 for the winning model. Winning gradient models where the gradient was less than 4.8mm were considered wins for the boundary model to ensure an underlying functional boundary blurred by the point spread function inherent to fMRI collection would not be considered a gradient (see Methods for details). It should also be noted that individualized axes of greatest change were determined using the boundary model, meaning the gradient models were fit on the axis that maximized the boundary model fit, thus steel-manning the boundary hypothesis.

Results largely mirrored the group level probabilistic shift results and individual contrast visualizations, showing strong evidence for the gradient hypothesis in preSMA, PMv, and aIns across visual and auditory contrasts, and for PMd in the visual contrast (Figure 8, Table 3, Supplemental Figure 6). In the visual modality, we examined 5 frontal lobe ROIs that exhibited COM shifts in either hemisphere. Strong evidence supported the gradient model over the boundary model in bilateral preSMA (77% vs. 11%), bilateral PMv (83% vs 6%), left aIns (63% vs. 25%), and left PMd (61% vs. 6%), all with gradient lengths significantly greater than 4.8mm. In contrast, evidence favored the boundary model in left mid-DLPFC (33% vs. 53%) where gradient length was not significantly greater than 4.8mm. The right hemisphere ROIs for aIns, PMd, and mid-DLPFC did not pass the COM shift test. One potential limitation of this criterion is that it could reject an ROI if subject-level gradients are too heterogeneous in location to create a spatial difference between sensory and WM COMs at the probabilistic group level. To examine whether hemispheric differences might exist with respect to gradients, we also performed the same individual-subject level analysis on these three ROIs. RH aIns and RH PMd, which failed the COM test, yielded similarly positive results for individual-subject gradients as did their respective left hemisphere ROIs, with the gradient model winning 71% vs. 6% in RH aIns and 42% vs. 5% in RH PMd with significant gradient lengths of 10.3 mm and 7.7 mm, respectively. Conversely, RH mid-DLPFC results showed some evidence for gradients with a significant gradient length, although the ratio of gradient wins to boundary wins was small (39% vs. 22%). Thus, we do not observe strong evidence for hemispheric differences in the visual functional organization for aIns or PMd but cannot rule it out for mid-DLPFC. Mean gradient length spanned 6.9 mm to 10.5 mm across these ROIs (see Table 3).

**Figure 8.**
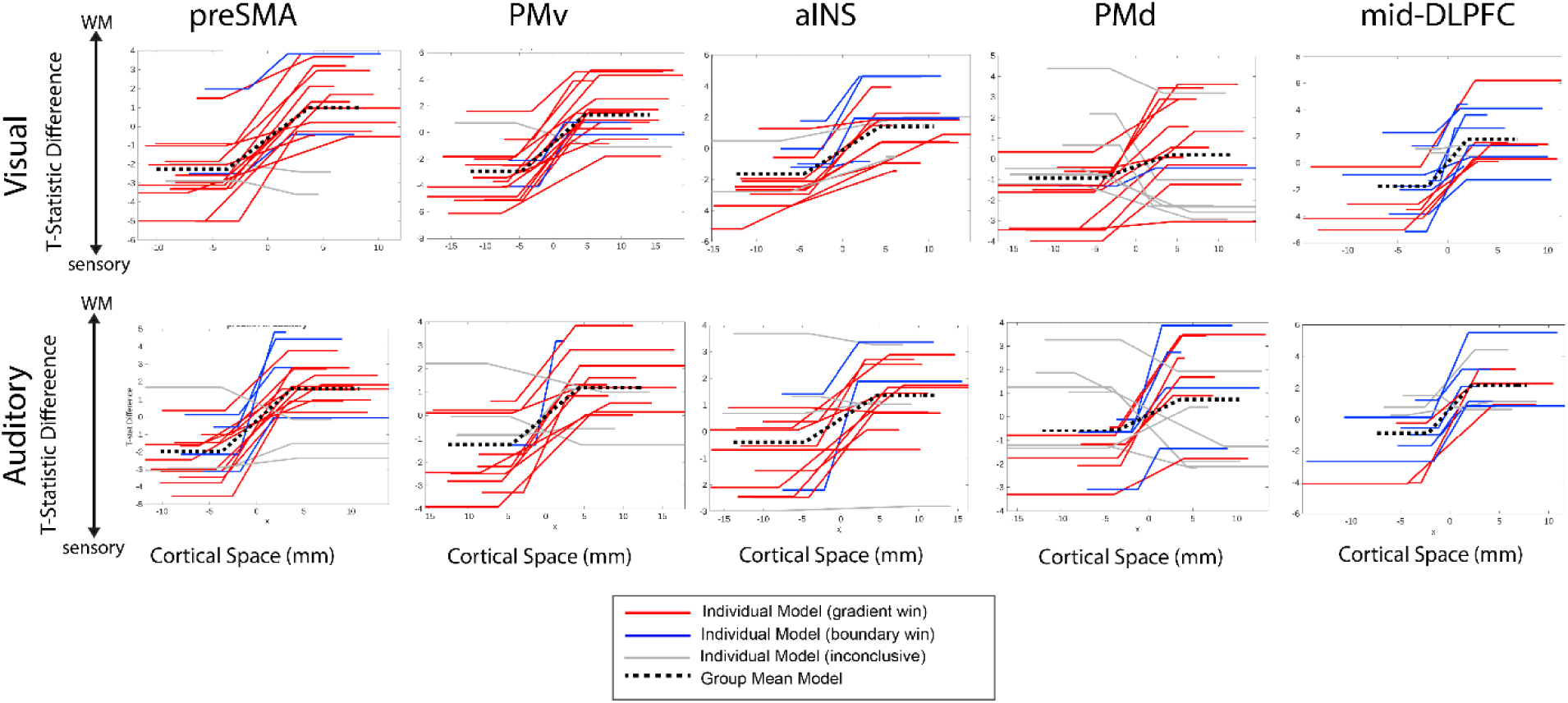
Mean slice model fits for each subject, ROI, and modality for the left hemisphere (see Supplemental Figure 6 for right hemisphere). All models’ x-values are aligned so that 0 corresponded to the best fit boundary point between sensory and WM activation for that subject/ROI. Red lines indicate gradient wins, blue indicates boundary wins, and grey indicates inconclusiveness. Group mean models are indicated by black dashed lines.

**Table 3.**
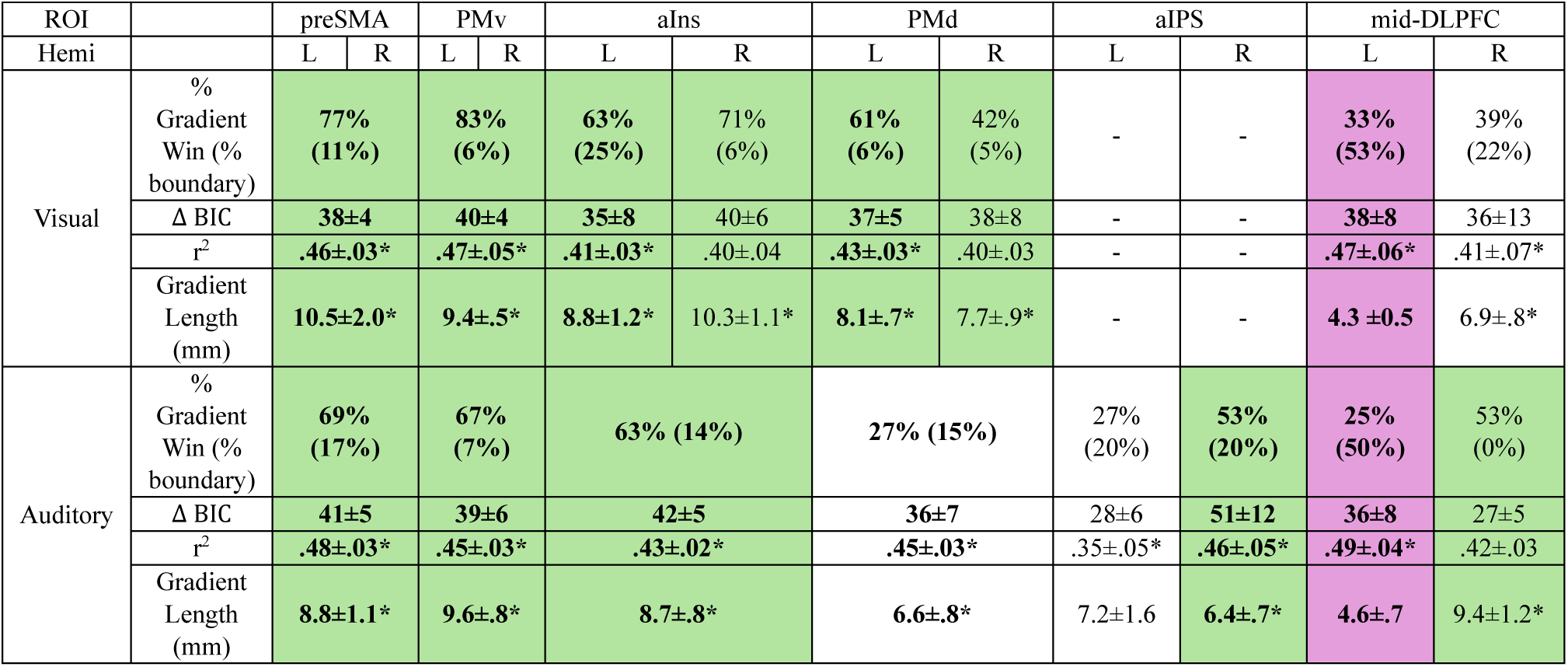
Comparison of gradient and sharp boundary models for visual and auditory working memory. Each modality was examined separately with ROIs defined for each subject. Gradient and boundary models were contrasted (see Methods, Fig 4). Green indicates ROIs in which the gradient model won at least twice as often as the boundary model and gradient length was significant. Magenta indicates ROIs in which the boundary model won more often than the gradient model. White indicates ROIs that were judged inconclusive. When only one hemispheric ROI passed the group-level COM shift test, the hemispheres were considered separately in that modality (bold text = passed COM shift test; plain text = failed COM shift test). The following information is displayed in sub-rows: % of model comparisons in which each model won, mean difference in BIC between gradient and boundary models, mean r-squared for the winningest model, and average length (mm) of the gradient portion across all fitted hinge functions. Means are displayed ± standard error. Asterisks indicate p<.05 corrected for multiple comparisons using FDR-BH against a null hypothesis of µ<.1 for r^2^ and µ<4.8mm for gradient length.

In the auditory modality, we examined 5 frontal lobe ROIs and 1 parietal lobe ROI that exhibited COM shifts in either hemisphere. Strong evidence supported the gradient model over the boundary model in bilateral preSMA (69% vs. 17%), bilateral PMv (67% vs 7%), bilateral aIns (63% vs. 14%), and right aIPS (61% vs. 6%), and evidence was weak in bilateral PMd (27% vs. 15%), all with significant gradient lengths. The boundary model was again favored in left mid-DLPFC (25% vs. 50%) (see Figure 8, Table 3) where gradient length was not significant. We also examined LH aIPS and RH mid-DLPFC, which failed the COM shift test. LH aIPS results were inconclusive (27% vs. 20%) with non-significant gradient length, and surprisingly RH mid-DLPFC supported the gradient model (53% vs. 0%) and had significant gradient length. Thus, we cannot rule out hemispheric difference in auditory WM organization in aIPS and/or mid-DLPFC. Mean gradient length spanned 6.4 mm to 9.6 mm across these ROIs (see Table 3).

Thus, overall, we present strong evidence that gradient-like models describe changes in sensory and WM activation better than boundary-like models in several areas of frontal cortex. PreSMA, PMv, and aIns exhibited strong evidence for gradients in both visual and auditory conditions. PMd gradients were observed in the visual modality but were inconclusive in the auditory modality. These gradients generally span 8-10mm of cortex (about 4 voxels) with sensory activation dominating rostrally and WM activation dominating caudally.

### Comparing sensory-working memory gradients with HCP-MMP1.0 boundaries

The Human Connectome Project Multi-Modal Parcellation 1.0 (HCP-MMP1.0 or MMP) is a widely used parcellation of the human cerebral cortex that leverages cortical architecture, task-based activation, resting state connectivity, and topography to delineate boundaries between functional areas (Glasser et al., 2016). One source the MMP draws on to inform boundaries is resting state functional connectivity (FC) gradients (i.e. changes in functional connectivity profiles across cortical space). But there appears to be little consideration of how sharp (steep and short) the FC gradient needs to be to indicate a boundary instead of a true gradient that spans significant cortex, indicating the lack of a clear boundary. Close examination of the visual WM (VWM) task (2-back) in the HCP/MMP dataset reveal multiple instances in frontal cortex where VWM activation straddles MMP parcel boundaries with FC gradients. This could potentially indicate a WM gradient across the parcellation “boundary” which was created partially using an FC gradient measure.

For example, the boundary between MMP parcels SCEF and 8BM follows a FC gradient that is spanned by VWM activation (Figure 9C), and the sensory-WM gradient identified here in preSMA also straddles the SCEF-8BM boundary (Figure 9 A, B). A similar motif appears again for both PMv and PMd. HCP/MMP reveals FC gradients between MMP ROIs 6a/FEF and i6-8 as well as between 8C and PEF while the VWM activation spans both of the boundaries created using the FC gradients. The sensory-WM gradient identified in PMv in our dataset partially straddles the 8C-PEF boundary as well, although there seems to be a mismatch between datasets in PMd activation. Nevertheless, HCP data seem to provide evidence supporting the hypothesis of meso-scale functional gradients in several frontal cortical regions, at least for the visual modality.

**Figure 9.**
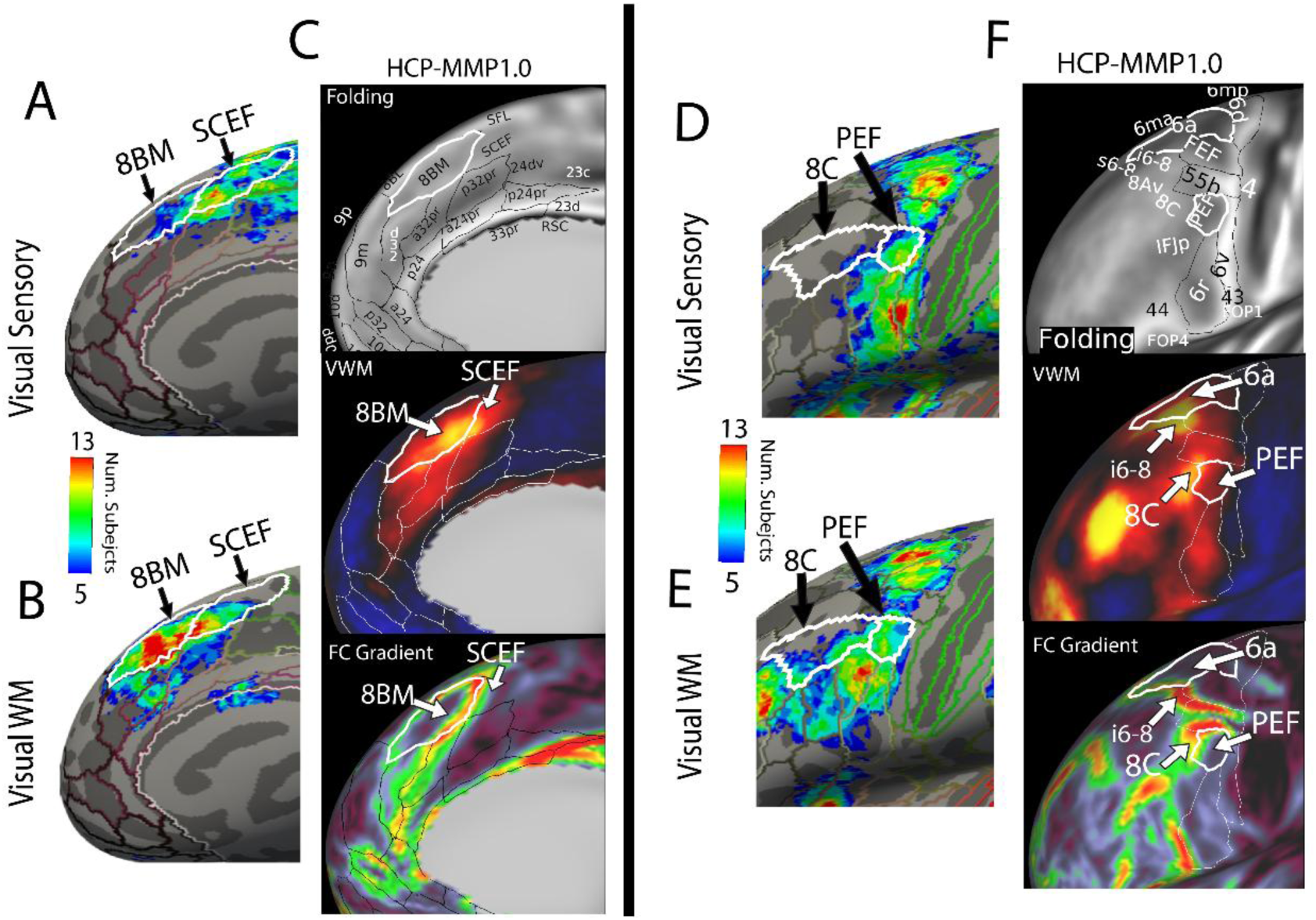
A) Probabilistic map of visual sensory activation with MMP parcel boundaries overlayed. SCEF and 8BM parcels are outlined in white and labeled. B) Probabilistic map of visual WM activation with MMP parcel boundaries overlayed. C) Showing frontal medial right hemisphere parcels (top), 2-back VWM group activation (middle), and group-level FC gradients (bottom). Data for panels & and F were downloaded from the BALSA Washington University – University of Minnesota Consortium of the HCP (https://balsa.wustl.edu/sceneFile/n889) under open access use terms. D), E), and F) are in the same format as A), B), and C) respectively, showing MMP parcels 8C, PEF, i6-8 and 6a.

### Domain general regions overlap working memory end of gradients

Assem et al. (2020) identified 7 “domain general” or “multiple demand” MMP frontal cortex parcels that consistently responded strongly over each of 3 cognitive task contrasts (math > story, WM 2-back > 0-back, and relational reasoning hard > easy). These regions are thought to engage in flexible cognitive control processes across task domain. We found that within our probabilistic ROIs, these domain general MMP parcels overlapped much more with our WM probabilistic maps (mean Dice coefficient of .42) compared to our sensory probabilistic maps (mean Dice coefficient of .17) for both modalities (Table 4, Supplemental Figure 7). This may suggest that the WM end of the sensory-WM gradients overlaps or feeds into domain general processing regions. Additionally, domain general parcels overlapped more with visual contrasts than auditory contrasts for both sensory (.23 vs. .10) and WM (.44 vs. .40), suggesting a modality asymmetry as previously reported (Noyce et al., 2017; Possidente et al., 2026; Tobyne et al., 2025).

**Table 4.** Dice coefficients expressing overlap between domain general MMP frontal cortex parcels and probabilistic maps for visual and auditory sensory and working memory contrasts in each ROI. Bolded values are from ROIs that were significant in the COM shift analysis.

| ROI | preSMA |  | PMv |  | aIns |  | PMd |  | mid-DLPFC |  |
| --- | --- | --- | --- | --- | --- | --- | --- | --- | --- | --- |
| Hemi | L | R | L | R | L | R | L | R | L | R |
| Visual Sensory | **.04** | **.19** | **.27** | **.25** | **.22** | .40 | **.06** | .19 | **.25** | .44 |
| Auditory Sensory | **.02** | **.05** | **.13** | **.07** | **.13** | **.12** | **.00** | **.11** | **.10** | .27 |
| Visual WM | **.35** | **.46** | **.47** | **.46** | **.45** | .45 | **.15** | .18 | **.79** | .44 |
| Auditory WM | **.30** | **.43** | **.40** | **.40** | **.33** | **.38** | **.10** | **.21** | **.75** | .68 |

## Discussion

In this study we sought to understand the meso-scale organization of the interaction between sensory processing and WM in frontal cortex by adjudicating between boundary and gradient hypotheses of functional organization. Our results demonstrated that bilateral preSMA, PMv, and aIns exhibit clear rostral-caudal sensory-WM gradients of function for both visual and auditory domains, whereas PMd exhibited strong gradient evidence primarily for the visual condition. Mid-DLPFC displayed hemispheric differences with LH evidence for a boundary between sensory and WM regions in both sensory modalities, while there was some evidence for gradients in RH. This may indicate that LH mid-DLPFC could be less involved in the integration and comparison of mnemonic and incoming sensory representations, and more involved with the maintenance and manipulation of mnemonic representations. The methods used accounted for smoothing incurred by the point spread function inherent in fMRI collection, fit gradient models on individualized axes that maximized the boundary model fit, and did not use smoothing during preprocessing. The gradients reported were typically in the range of 8-10 mm. By comparison, smaller frontal ROIs in the MMP parcellation can nearly span similar distances (ex. i6-8 spans about 11.5mm in the dorsal-ventral direction). This suggests the reported functional gradients could nearly span entire cortical regions or serve as extended transition zones between more selective regions.

Sensory WM is tasked with encoding external stimuli but must also store and manipulate information to support behavioral goals. If the influence of these bottom-up and top-down constraints are viewed as a continuum, one potential advantage of a gradient structure is permitting external modulation of balance between sensory and cognitive influences, depending on the current task demands. The overlap between domain-general parcels and the WM end of gradients presented here support this idea further. Many frontal cortical neurons exhibit mixed-selectivity for multiple factors of WM and cognitive control (Dang et al., 2022; Rao et al., 1997; Rigotti et al., 2013), offering computational advantages in the diversity of possible input-output functions (Rigotti et al., 2013; Tye et al., 2024). The meso-scale organization of mixed-selectivity neurons is not yet clear, but we suggest functional gradients offer an efficient means of organizing and modulating cognitive information in frontal cortex. This suggests a possible resolution to the dual-representation problem: frontal gradients could hold sensory and WM representations separately at either end, while permitting coordination between them in the middle. This interpretation, however, depends on stimulus-based representations in these areas, something that would require stimulus feature decoding to show definitively, which this task paradigm does not support.

The notion of gradients of cortical organization challenges long-held notions of anatomically and functionally distinct cortical zones (Brodmann, 1909; Vogt & Vogt, 1919). Bailey & von Bonin (1951) argued that the notion of sharp cytoarchitectonic boundaries within cortex was greatly overstated. Modern architectonic analysis provides more nuance, noting that areal boundaries vary from abrupt to more gradual changes (Amunts & Zilles, 2015). Notably, recent analysis of human premotor cortex reports anatomical boundaries between regions along the dorsal-ventral axis, but more gradual changes along the rostral-caudal axis (Ruland et al., 2025). This appears consistent with the location and orientation of premotor cortical sensory-WM gradients reported here. Neuroimaging research still largely adheres to a sharp boundary notion of cortical areas (Buckner et al., 2011; Glasser et al., 2016; Schaefer et al., 2018) even as evidence builds for smaller and potentially overlapping functional zones (Assem et al., 2025; DiNicola & Buckner, 2026; Somers et al., 2021). Algorithmic approaches that assume the existence of boundaries will necessarily draw such boundaries, even along a true gradient. The MMP (Glasser et al., 2016) relies partly on measurements of change in intrinsic FC over cortical space to define some cortical boundaries, overlooking the possibility of true functional gradients. We note here multiple instances from that dataset in which VWM activation in frontal cortex spans neighboring parcels denoted by strong gradients in FC. These appear to coincide with the sensory-WM gradients we observe in preSMA and PMv. More broadly, we suggest that next-generation cortical parcellation schemes should incorporate the potential for gradient zones.

Conceptualizing these frontal sensory-WM regions as gradients instead of boundaries has important implications for neuroimaging analysis methodology, interpretation of results, and clinically relevant hypothesis generation. For example, WM analysis using statistically thresholded ROIs on activity maps is likely to have ROIs contaminated with sensory-drive activated regions, and smoothing will exacerbate this problem. Additionally, the ability to identify and measure these gradients could provide clinically useful features for disorders where interaction between sensory and WM processing is thought to be dysfunctional, like ASD (Alsaedi, 2025; Pastor-Cerezuela et al., 2020; Stevenson et al., 2021), schizophrenia (Chen et al., 2009; Hamilton, et al., 2018; Javitt & Freedman, 2014), ADHD (Kofler et al., 2020; Li et al., 2023), and dyslexia (Beneventi et al., 2010; Fostick & Revah, 2018; Laasonen et al., 2012).

Although all five frontal ROIs exhibited evidence for gradients in at least one hemisphere and/or one sensory modality, it seems likely that there are important differences in the information coded along these gradients. Prior work has demonstrated distinct zones within PMd, PMv, preSMA, and mid-DLPFC that are biased toward visual or auditory working memory (Assem et al., 2022; Michalka et al., 2015; Noyce et al., 2017; Tobyne et al., 2025) (see Supplemental Figure 8 for probabilistic maps from the present data). For instance, PMd contains both a visual-biased region in the superior precentral sulcus (sPCS) and an auditory-biased region in the transverse gyrus intersecting precentral sulcus (tgPCS) or area 55b. These finer sensory-biased subzones may explain the modality differences observed in PMd.

Our analyses were successful in revealing several sensory-WM functional gradients within frontal cortex, however, our search was limited to relatively large areas that exhibited group-level differences. It may be the case that some rejected ROIs also display boundary and/or gradient properties that were not evaluated due to insufficient power to detect group level spatial differences. Additionally, we could have overlooked regions that may exhibit finer scale differences in patterns of stimulus drive and WM, especially if they are more heterogeneous across individuals. A gradient zone may exist as an isolated, self-contained region or it might terminate at one or both ends in regions that exhibit strong selectivity for one type of information. The center-of-mass shift criterion that we applied to the group-level data would be more likely to detect the latter type of gradient and potentially miss some isolated gradient zones. Finer spatial scale imaging, particularly at 7 Tesla, could also be beneficial due to the smaller point spread function (Shmuel et al., 2007).

Further work should explore these gradients in the context of different stimulus types and across the subcomponents of WM. This analysis is limited by the nature of the experimental paradigm, which only presented basic visual and auditory stimuli. Although more complex stimuli used in other work have successfully identified sensory and WM activation in similar cortical regions (Noyce et al., 2022), gradient vs. boundary analysis was not performed for these datasets. Given the possibility of stimulus category-biased patches surrounding domain-general frontal regions suggested by Assem et al. (2025), it seems plausible that sensory-WM gradients may shift in a stimulus-dependent manner. Additionally, using a long-delay paradigm instead of the short delay employed here would enable analysis of encoding, maintenance, and retrieval components of WM, and whether they are differentially related to sensory-WM gradients found here.

This investigation adds to the literature (Farrugia et al., 2024; Lefco et al., 2020; Shen et al., 2023; Tian & Zalesky, 2018) advocating that the meso-scale functional organization of some cortical regions are better described as gradients than boundaries. Some of these gradient reports derive primarily from analyses of functional connectivity changes, leaving the cognitive/informational axes incompletely specified. We do not mean to suggest cortical gradients are limited to sensory-WM axes, but rather that gradients could be an efficient organizational scheme to handle conditions in which cognitive control demands substantial flexibility. It is also worth noting that multiple gradients, along different principal axes, could be overlaid within a single region as a means of supporting multi-dimensional cognitive control.

## Data Availability

Unprocessed task and structural MRI data are publicly available on OpenNeuro at doi:10.18112/openneuro.ds007231.v1.1.1.

## Code Availability

Custom Matlab code used to produce this work can be found at https://github.com/fmri/Frontal_Gradients_Boundaries/.

## CRediT Author Statement

**Thomas Possidente**: Conceptualization, Formal analysis, Methodology, Software, Validation, Visualization, Writing – original draft, Writing - review & editing, **Vaibhav Tripathi:** Data curation, Investigation, Writing – review & editing, **Sangil Lee:** Conceptualization, Methodology, Writing – review & editing**, David Somers:** Conceptualization, Funding acquisition, Methodology, Supervision, Project Administration, Writing – original draft, Writing – review & editing.

## Acknowledgments

This work is funded by National Science Foundation grant BCS-1829394 to D.C.S. This work involved the use of instrumentation supported by the NSF Major Research Instrumentation grant BCS-1625552. Data was analyzed on a high-performance computing cluster supported by the ONR grant N00014-17-1-2304.

Data for Figure 9 were provided by the Human Connectome Project, WU-Minn Consortium (Principal Investigators: David Van Essen and Kamil Ugurbil; 1U54MH091657) funded by the 16 NIH Institutes and Centers that support the NIH Blueprint for Neuroscience Research; and by the McDonnell Center for Systems Neuroscience at Washington University.

We thank Dr. Abigail Noyce for auditory stimuli, and Dr. David Beeler, Dr. Ryan Marshall, Dr. Stephanie McMains, and Shruthi Chakrapani for scanning assistance. We acknowledge the University of Minnesota Center for Magnetic Resonance Research for use of the multiband-EPI pulse sequences.

