## Supplemental Figures/Tables for "Gradients of function between sensory drive and working memory in human frontal cortex"

| Equation | Parameter | Initializing Estimate |
|---|---|---|
| Boundary (eq 1) | $x_1$ (step location x) | Mean of x coordinates in ROI |
| Boundary (eq 1) | $x_2$ (maximum y) | Mean of positive group-level T-statistics |
| Boundary (eq 1) | $x_3$ (minimum y) | Mean of negative group-level T-statistics |
| Gradient (eq 2) | $x_1$ (x start of hinge) | $25^{th}$ percentile of x coordinates |
| Gradient (eq 2) | $x_2$ (x end of hinge) | $50^{th}$ percentile of x coordinates |
| Gradient (eq 2) | a (minimum y) | Mean of negative group-level T-statistics |
| Gradient (eq 2) | b (maximum y) | Mean of positive group-level T-statistics |

Supplemental Table 1 | Estimates to initialize each parameter when fitting boundary and gradient equations.

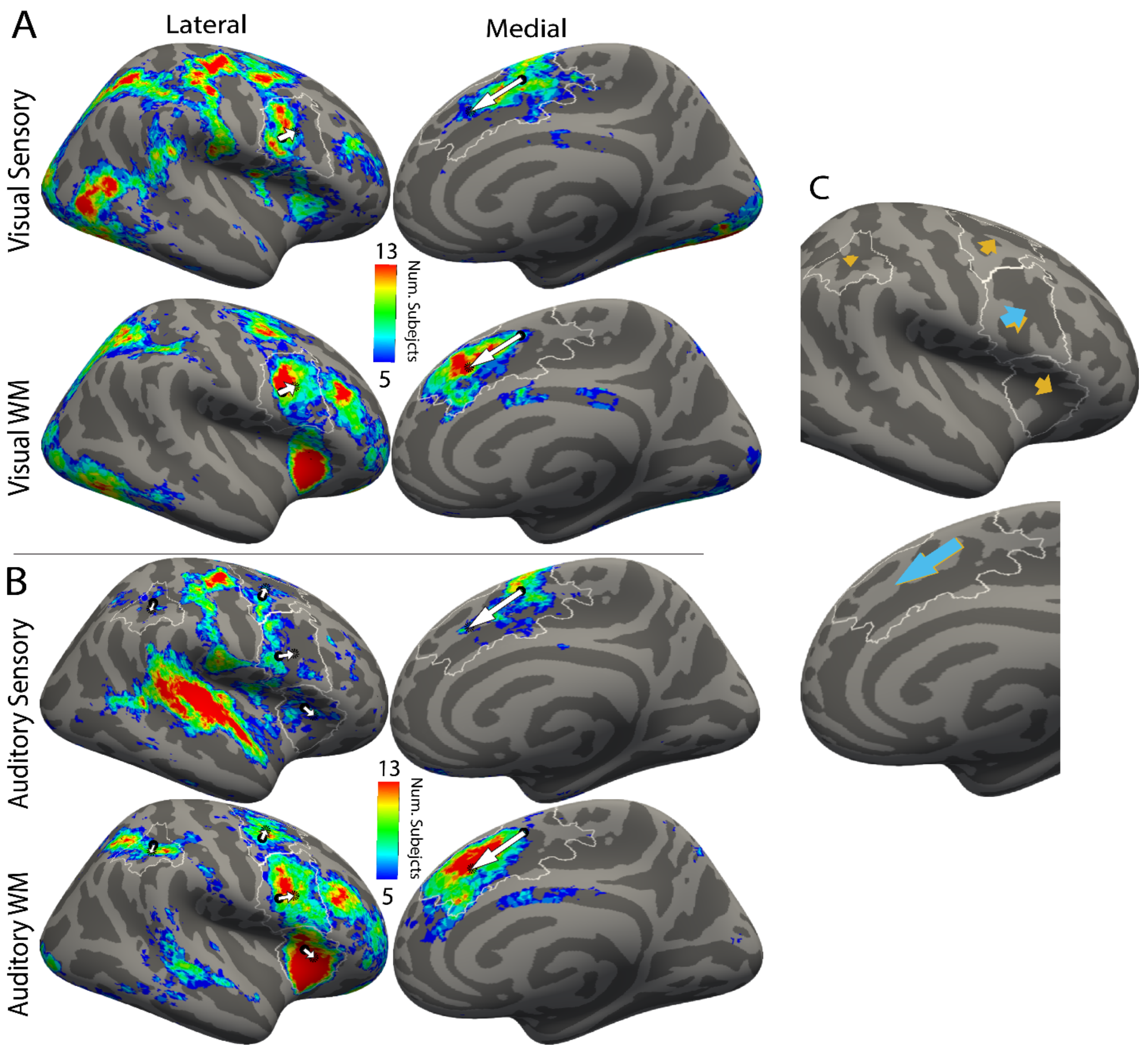


Supplemental Figure 1 | A) Probabilistic maps of visual sensory activation (top) and visual WM (bottom) with lateral (left) and medial (right) views. White arrows represent change of COM location from sensory to WM within each ROI. Black circles represent COMs for the map being displayed while black dotted circles represent COMs for the opposite gradient contrast. White borders show the probabilistic search spaces for each ROI. B) Same as A except using auditory sensory activation (top) and auditory WM (bottom). C) Visual (blue) and auditory (orange) sensory-WM gradient directions consolidated from A and B and enlarged. Lef hemisphere results are shown in Figure 1.

| ROI | Hemisphere | Visual | Auditory |
|---|---|---|---|
| preSMA | L | **85%** | **90%** |
| | R | **90%** | **85%** |
| PMv | L | **90%** | **75%** |
| | R | **85%** | **75%** |
| aINS | L | **80%** | **80%** |
| | R | 85% | **95%** |
| PMd | L | **90%** | **80%** |
| | R | 95% | **85%** |
| mid-DLPFC | L | **75%** | **60%** |
| | R | 90% | 75% |
| aIPS | L | - | 75% |
| | R | - | **75%** |

Supplemental Table 2 | Percent of ROIs identified in individual subjects for each modality and hemisphere in which there was a significant spatial difference in WM and sensory COMs. Bolded values are from ROIs that were significant in the COM shift analysis.

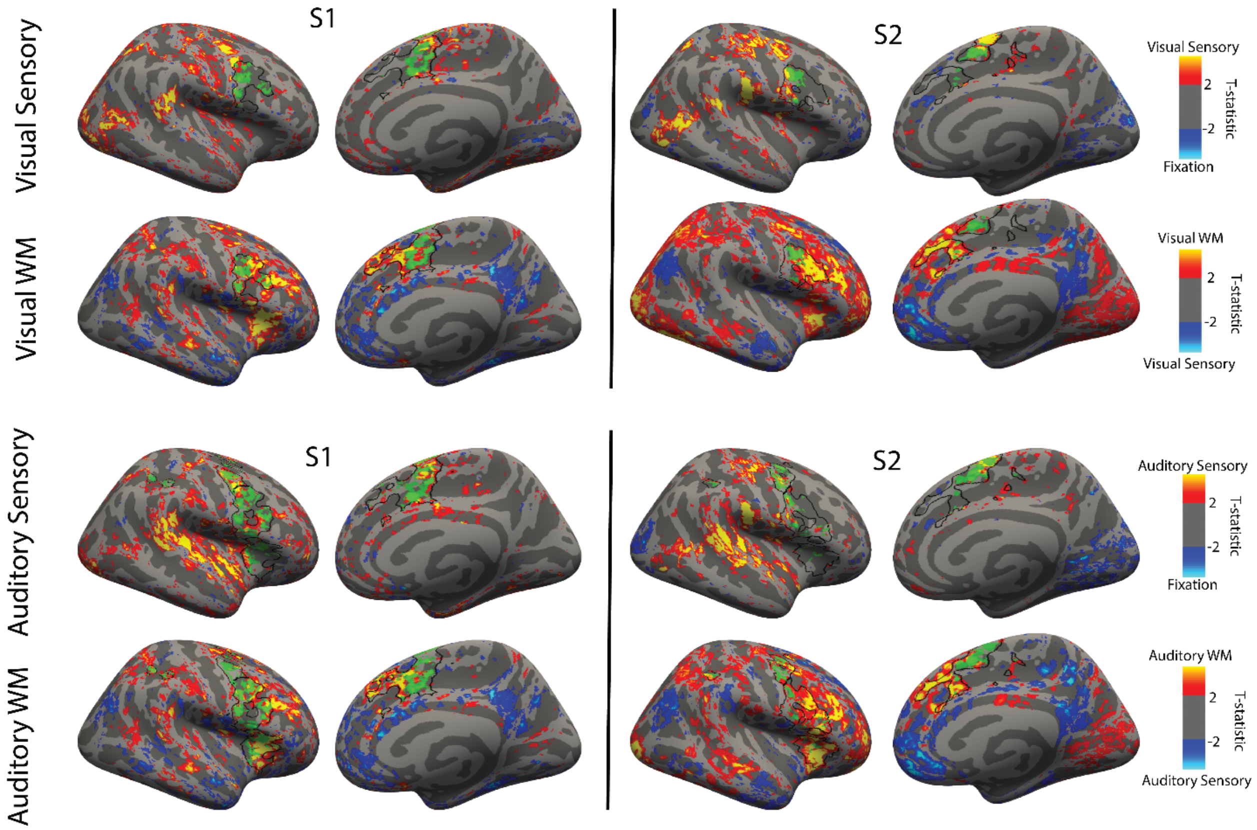


Supplemental Figure 2 | Arrangement is analogous to Figure _A, but instead of probabilistic maps, two example subjects' T-statistic maps are shown using a T>2 threshold. Black outlines show individual-specific ROIs (identified using WM and sensory vs fixation contrast for each modality separately). ROIs that did not pass the inclusion criteria are not outlined in black. Green patches represent areas of overlap between the WM and sensory T-statistic maps. The left hemisphere version of this figure is shown in Figure 6.

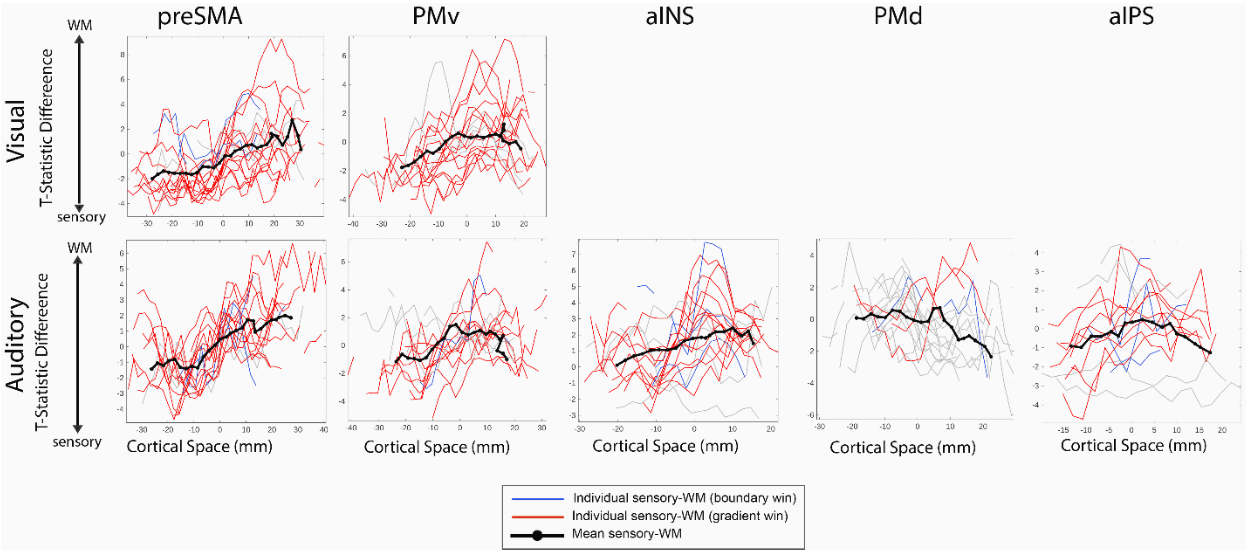


Supplemental Figure 3 | Binned T-statistics difference plotted across each ROI for each individual subject (visual and auditory). Thick black/gray lines show the mean of sensory/WM contrasts across subjects. Mean lines are only displayed if there are at least 5 subjects to average across. Only the right hemisphere results are shown here (see Figure 7 for left hemisphere).

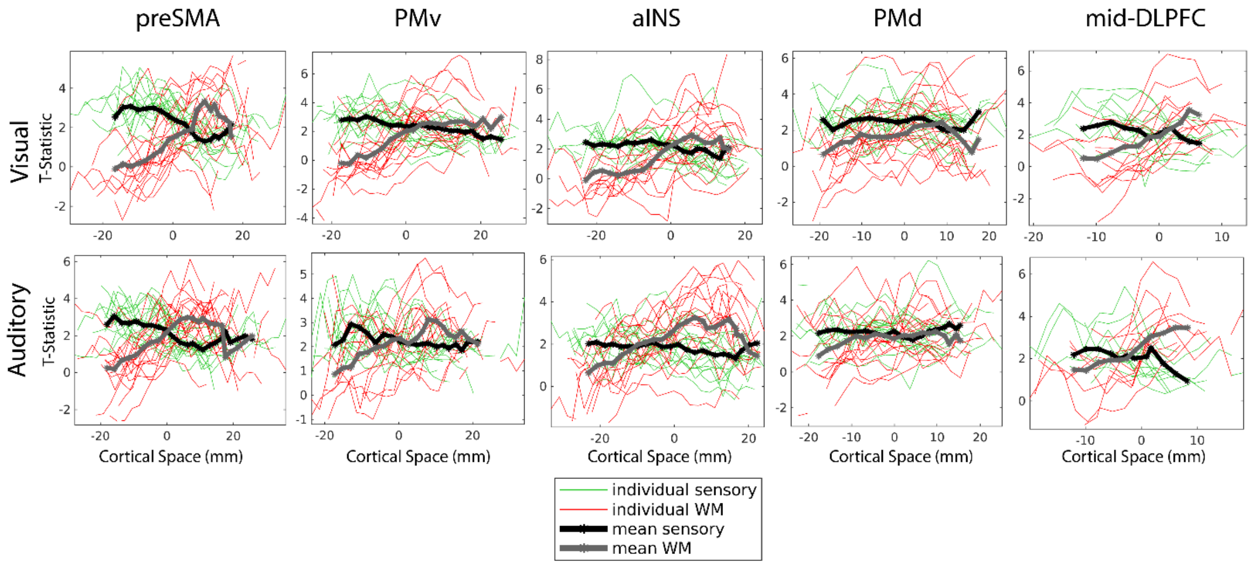


Supplemental Figure 4 | Binned T-statistics plotted across each ROI for each individual subject (visual and auditory) with sensory contrasts in green and WM contrasts in red. Thick black/gray lines show the mean of sensory/WM contrasts across subjects. Mean lines are only displayed if there are at least 5 subjects to average across. Only the left hemisphere results are shown here (see Supplemental Figure 5 for right hemisphere).

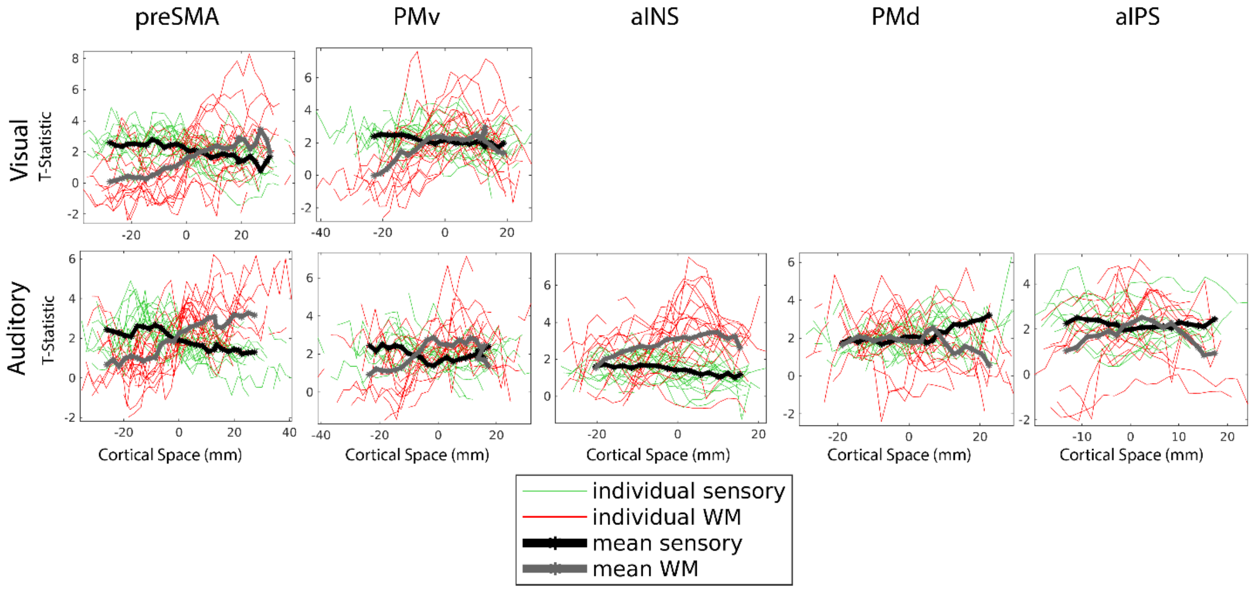


Supplemental Figure 5 | Binned T-statistics plotted across each ROI for each individual subject (visual and auditory) with sensory contrasts in green and WM contrasts in red. Thick black/gray lines show the mean of sensory/WM contrasts across subjects. Mean lines are only displayed if there are at least 5 subjects to average across. Only the right hemisphere results are shown here (see Supplemental Figure 4 for left hemisphere).

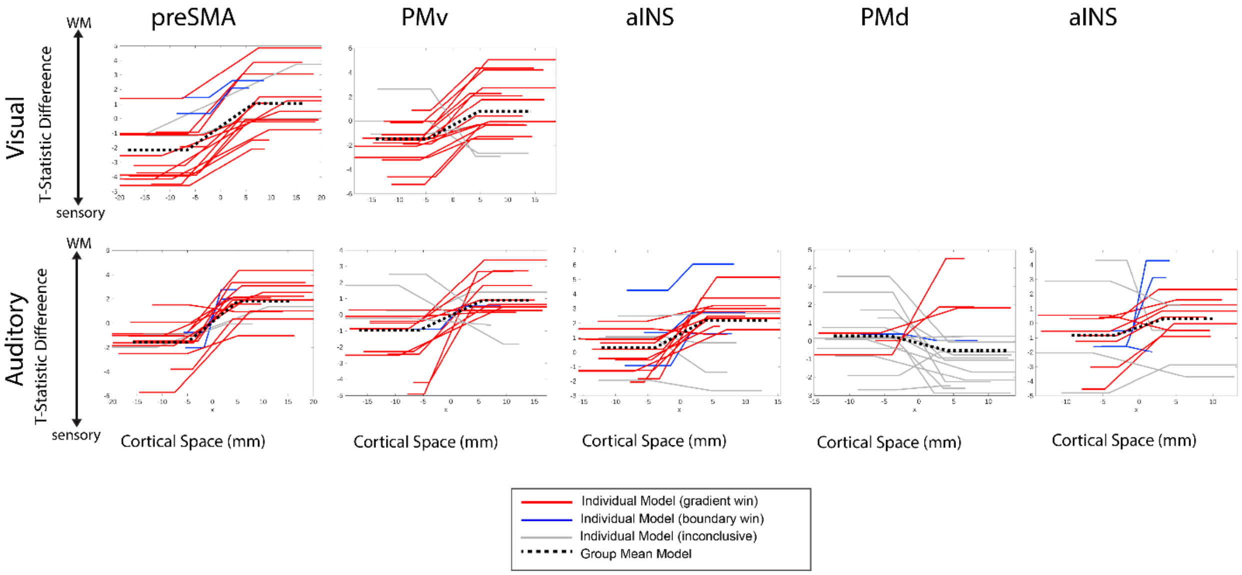


Supplemental Figure 6 | Mean slice model fits for each subject, ROI, and modality for the right hemisphere (see Figure 8 for left hemisphere). All models' x-values are aligned so that 0 corresponded to the best fit boundary point between sensory and WM activation for that subject/ROI. Red lines indicate gradient wins, blue indicates boundary wins, and grey indicates inconclusiveness. Group mean models are indicated by black dashed lines.

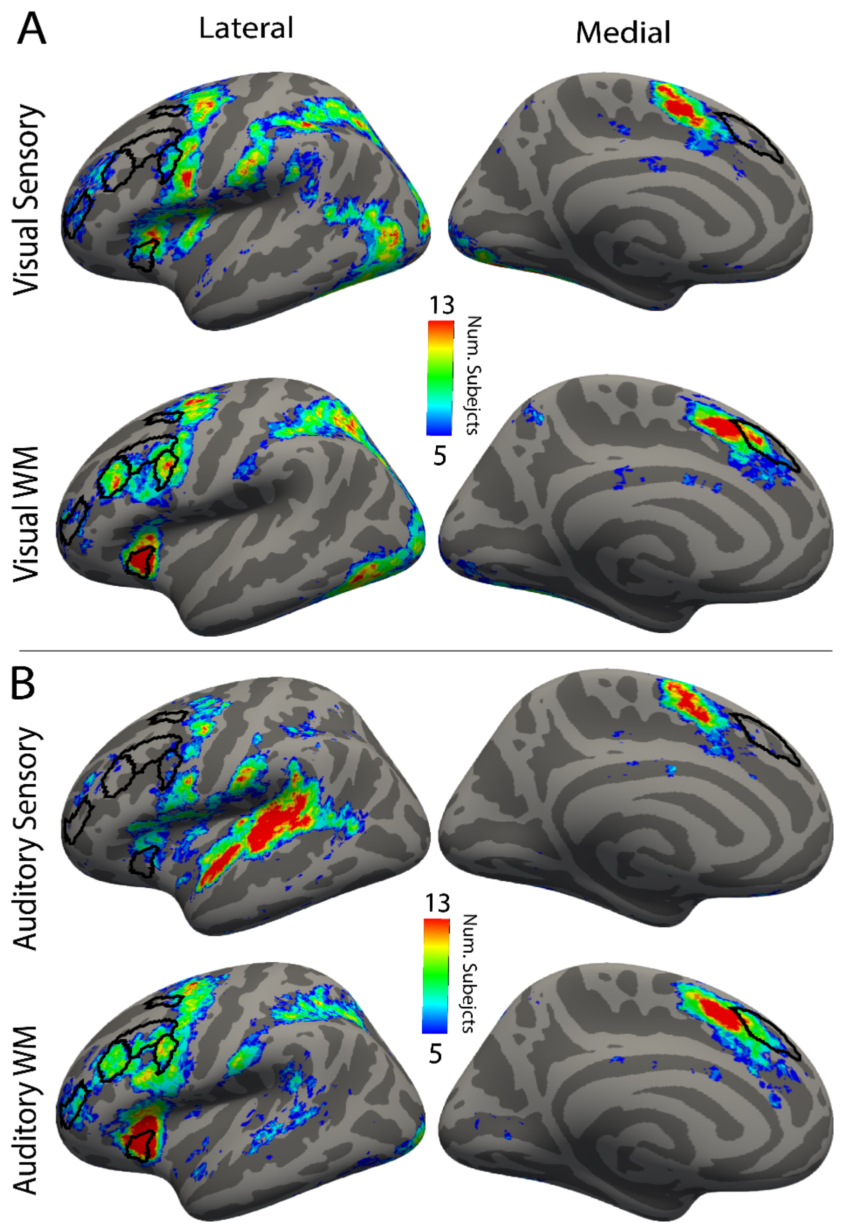


Supplemental Figure 7 | Probabilistic maps for visual and auditory sensory and WM activation are displayed in the same manner as Figure 5 and Supplemental Figure 1. Black outlines show domain general parcels from the MMP.

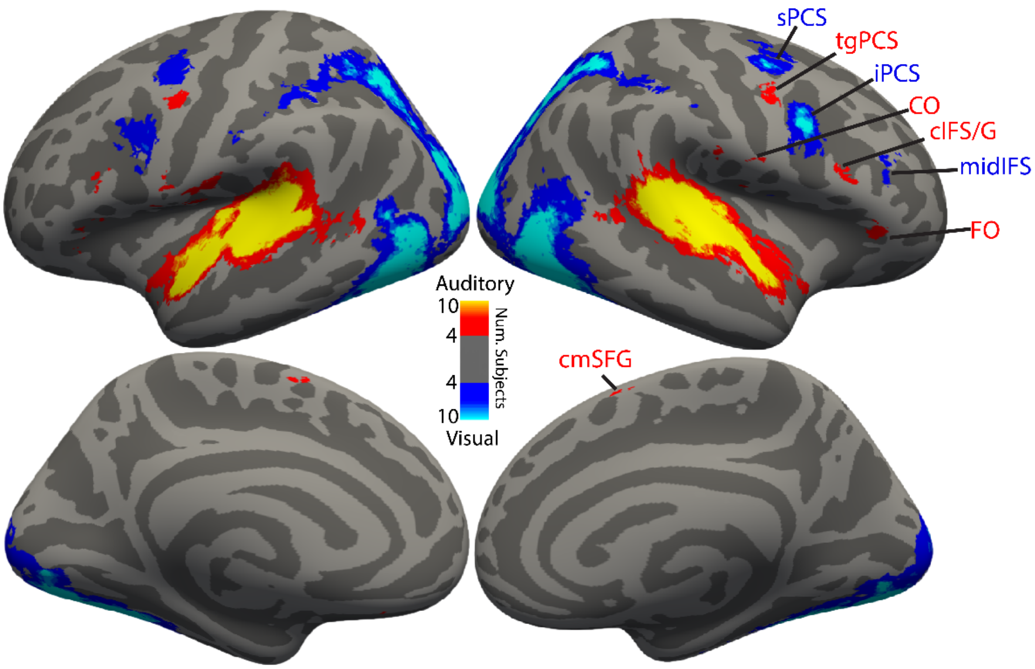


Supplemental Figure 8 | Probabilistic map showing sensory-bias regions. Maps were created by subtracting individual subject *visual WM + SMC > Fixation* t-statistic maps from *auditory WM + SMC > Fixation* t-statistics maps, then thresholding the resulting map at t > 2 and t < -2 to create an auditory-bias mask and visual-bias mask respectively. These masks were added together for each subject and the resulting visual-bias probabilistic was subtracted from the resulting auditory-bias probabilistic to create the sensory-bias probabilistic which was then thresholded at N >= 4 (20%) subjects.
